# Microfluidic single-cell scale-down bioreactors: A proof-of-concept for the growth of *Corynebacterium glutamicum* at oscillating pH values

**DOI:** 10.1101/2021.12.30.474512

**Authors:** Sarah Täuber, Luisa Blöbaum, Valentin Steier, Marco Oldiges, Alexander Grünberger

**Author notes:** **Correspondence** (A. Grünberger).

## Abstract

In large-scale bioreactors, gradients in cultivation parameters such as oxygen, substrate and pH result in fluctuating environments. pH fluctuations were identified as a critical parameter for bioprocess performance. Traditionally, scale-down systems at the laboratory scale are used to analyze the effects of fluctuating pH values on strain and thus process performance. Here, we demonstrate the application of dynamic microfluidic single-cell cultivation (dMSCC) as a novel scale-down system for the characterization of *Corynebacterium glutamicum* growth using oscillating pH conditions as a model parameter. A detailed comparison between two-compartment reactor (two-CR) scale-down experiments and dMSCC was performed for one specific pH oscillation between reference pH 7 (∼ 8 min) and disturbed pH 6 (∼2 min). Similar reductions in growth rates were observed in both systems (dMSCC 21% and two-CR 27%). Afterward, systematic experiments at different symmetric and asymmetric pH oscillations between pH ranges of 4–6 and 8–11 and different intervals from 1 minute to 20 minutes, were performed to demonstrate the unique application range and throughput of the dMSCC system. Finally, the strength of the dMSCC application was demonstrated by mimicking fluctuating environmental conditions within large-scale bioprocesses, which is difficult to conduct using two-CRs.

## 1. Introduction

In industrial bioreactors, the production host is subject to fluctuations environmental conditions, e.g., oxygen supply, nutrients and other process parameters that are negligible in laboratory-scale bioreactors, but can have a decisive influence on metabolism, yield and productivity (Nadal-Rey et al., 2020). These gradients, which result in fluctuating environments, are created by insufficient mixing in large volumes (Crater & Lievense, 2018) and can affect the cell over a time range of a few seconds to several minutes depending on the size of the reactor (Käß, Junne, Neubauer, Wiechert, & Oldiges, 2014).

In addition to oxygen and the main nutrient concentration, pH was identified as a critical parameter for the success of industrial bioprocesses (Klask, Kliem-Kuster, Molitor, & Angenent, 2020; Pohlscheidt et al., 2009). Typically, pH is controlled during cultivation to maintain optimal conditions for growth and productivity (Champagne, Gardner, & Doyon, 1989). Due to pH control and imperfect mixing, pH inhomogeneities can have three origins: chemical, biological and physical (Cortés, Flores, Bolívar, Lara, & Ramírez, 2016). The chemical origin is the selective addition of acid and base during pH control, which results in different pH zones (Amanullah, McFarlane, Emery, & Nienow, 2001; Lara, Galindo, Ramírez, & Palomares, 2006; Takors, 2012). Biological causes include the metabolic activity of the microorganisms. Nutrients such as glucose can be consumed in large quantities while consuming high amounts of oxygen (Limberg, Joachim, Klein, Wiechert, & Oldiges, 2017) to maintain metabolic balance and high consumption rates of carbon sources. These zones with surplus substrate and limited oxygen favor the formation of acidic byproducts, which may influence the pH. Many biotechnological products are weak acids such as acetate, lactate, some amino acids or ascorbic acid (Sahm, Antranikian, Stahmann, & Takors, 2013). As a result of chemical and biological factors, pH gradients for microbial processes are expected to range from pH 4 in the bulk liquid to pH 9 near the base addition zone (Lara et al., 2006; Reuss, Schmalzriedt, & Jenne, 1994). When cells are exposed to the extreme pH values near the addition point of the pH control, their viability and physiology might be affected (Hansen et al., 2016). Physical origins include hydrostatic pressure differences in large reactors that lead to increased O_2_/CO_2_ solubility at the bottom (Bothun, Knutson, Berberich, Strobel, & Nokes, 2004; Lopes, Belo, & Mota, 2014), the latter being a source of pH variation.

Currently, scale-down bioreactors are the system of choice for analyzing the influence of bioreactor inhomogeneity and the resulting gradients on the process performance of production strains during scale-up (Neubauer & Junne, 2016). Typically, single vessel, multicompartment or segmented stirred bioreactors are used (Lara et al., 2006; Neubauer & Junne, 2010). Here, different reactor types such as stirred-tank reactors (STRs) and plug flow reactors (PFRs) can be combined in two-compartment or more setups in various combinations (Limberg, Pooth, Wiechert, & Oldiges, 2016), where at least one STR compartment is used to mimic the bulk reactor zone. For example, scale-down bioreactors can mimic the bioreactor inhomogeneity of industrial-scale bioreactors (> 1 m^3^) by changing or oscillating the agitation rate, gassing conditions and substrate concentration by pulsating or oscillating substrate feed (Crater & Lievense, 2018). Scale-down systems are often used to analyze gradients of two or three parameters, e.g., substrate and dissolved oxygen (DO) concentration (Limberg et al., 2016) or substrate, DO and pH value (Olughu, Nienow, Hewitt, & Rielly, 2020). Limberg et al. used two-compartments, STR-PFR and STR-STR, for the analysis of byproduct formation in *Corynebacterium glutamicum*. A reduced growth rate and an increase in the intermediary formation of byproducts (L-lactate and L-glutamate) due to the gradients were found compared to a single reactor reference cultivation. The byproduct formation differed between the scale-down setups (i.e., STR-STR vs. STR-PFR). Recent scale-down studies have also identified pH fluctuations as critical and to date slightly neglected scale-up parameter, because it seems to imperil the robustness of microorganisms against other scale-up parameters such as oxygen (Limberg et al., 2017). Currently, the influence of pH gradients is highly disputed. Some researchers have reported from computational fluid dynamics (CFD) simulations, that the effects of fluctuations are very small or only occur in combination with fluctuations of other process parameters (Spann et al., 2019), while other researchers have experimentally observed a significant influence on the overall growth and productivity of a bioprocess (Amanullah et al., 2001; Cortés et al., 2016; Langheinrich & Nienow, 1999; Onyeaka, Nienow, & Hewitt, 2003; Spann et al., 2019).

A detailed analysis of the effects of fluctuations and gradients requires a system that enables the precise manipulation of environmental conditions to study physiological reactions and gene expression under fluctuating conditions. For this purpose, some microfluidic devices have already been developed that allow dynamic environmental conditions by combining valves or even pumps with the microfluidic system (Kaiser et al., 2018; Uhlendorf et al., 2012). One of these systems is the dMSCC (dynamic microfluidic single-cell cultivation), which allows specific and precise investigation of fluctuations (Täuber, Golze, Ho, von Lieres, & Grünberger, 2020). dMSCC can precisely control environmental conditions in time frames between 5 seconds and hours. This system can be used to create pH profiles with two or three different pH values with a high temporal oscillation frequency and thus enable a wide range of experiments with pulse frequency modulation, pulse width modulation and pulse amplitude modulation (Täuber, von Lieres, & Grünberger, 2020). In combination with live-cell imaging, this system allows the analysis of cellular response with high spatiotemporal resolution to study the behavior of populations at the single-cell level based on specific medium oscillations, for example, between BHI and PBS as shown by Täuber et al. (Täuber, Golze, et al., 2020).

In this study, the dMSCC system was applied for the first time for a detailed and systematic analysis of the effect of pH oscillation on the growth behavior of *C. glutamicum. C. glutamicum* is one of the most important platform organisms for the large-scale production of amino acids, e.g., L-lysine (Lee, Na, Kim, Lee, & Kim, 2016) and L-glutamate (Becker & Wittmann, 2017) and other industrially relevant bulk and fine chemicals (Becker, Kuhl, Kohlstedt, Starck, & Wittmann, 2018). First, we demonstrated the application of dMSCC as a new scale-down system for characterizing the growth of *C. glutamicum* under oscillating pH conditions. Based on traditional scale-down experiments with two-compartment reactor (two-CR), a detailed comparison with dMSCC was performed for a pH fluctuation between pH 6 and pH 7. Afterward, dMSCC was used for a systematic analysis of the growth behavior of *C. glutamicum* exposed to different oscillations between pH 7 and several pH values (pH 4–11) at various total interval durations *T* from 1 min to 20 min. Finally, we demonstrated for the first-time fluctuating pH environmental conditions from a large-scale bioprocess.

## 2. Materials and Methods

### 2.1 Medium preparation, bacterial strain and preculture

All cultivations were performed in CGXII minimal medium (Unthan et al., 2014) containing the following per liter of demineralized water: 20 g of (NH_4_)_2_SO_4_, 1 g of K_2_HPO_4_, 1 g of KH_2_PO_4_, 5 g of urea, 13.25 mg of CaCl_2_·H_2_O, 0.25 g of MgSO_4_·7H_2_O, 10 mg of FeSO_4_·7H_2_O, 10 mg of MnSO_4_·H_2_O, 0.02 mg of NiCl_2_·6H_2_O, 0.313 mg of CuSO_4_·5H_2_O, 1 mg of ZnSO_4_·7H_2_O, 0.2 of mg biotin and 40 g of D-glucose. The concentration of protocatechuic acid (PCA) was 30 mg/L in standard medium. In selected experiments (chapters 3.1 and 3.3) PCA was replaced by citrate at a final concentration of 3.84 mg/L. For shaking flask cultivation 42 g/L MOPS buffer was added. The solutions were autoclaved or sterile filtered. All used chemicals were purchased from Carl Roth. Prior to each dMSCC experiment, the medium was prepared, and the pH was adjusted with NaOH or H_3_PO_4_ before starting the experiments. The medium was additionally sterile filtered to prevent channel clogging during the experiments.

dMSCC osmolarity control experiments (see chapter 3.1) were performed with a higher salt concentration. Therefore, the concentration of NaCl was adjusted to 18.7 g/L resulting in an osmolarity of ∼ 1340 mOsmol/kg.

In this study, *C. glutamicum* ATCC 13032 was used. In the preculture, *C. glutamicum* was cultivated in CGXII medium at 30 °C in a rotary shaker at 120 rpm (Ecotron, Infors, Germany). The preculture was inoculated from a glycerol stock into a 100 mL shaking flask with 10 mL working volume and cultivated overnight. The main culture was inoculated from the preculture with a starting optical density (OD_600_) of ∼ 0.05.

### 2.2 Two-CR scale-down cultivations

Laboratory scale cultivations were performed in a scale-down bioreactor configuration in a fourfold parallel bioreactor setup. Two STRs were connected in an STR-STR scale-down bioreactor setup as described by Limberg et al. (Limberg et al., 2016), and experiments were performed in biological duplicates. The volumetric distribution of the initial working volume of 1000 mL between the two STR compartments under scale-down operation was set to 20% in the smaller compartment (STR2) (Fig. 1). DO and pH were individually controlled in both compartments. pH control was achieved through the addition of HCl (6 M) or NaOH (6 M). DO was controlled at 30% through a cascade involving the stirrer speed, aeration rate, and oxygen concentration in inlet air. Cultivations were carried out in batch mode at 30 °C and pH = 7.0. For cultivations under pH oscillation, the set point in STR2 was pH = 6.0 after the onset of exponential growth was observed. Biomass measurements were conducted using optical density measurements at 600 nm. According to the calculation published by Limberg et al. (Limberg et al., 2016), the average residence time of cells in STR2 was calculated to be between 2 and 2.5 minutes. Precultures were carried out in baffled 500 mL shake flasks containing 50 mL of CGXII medium (Limberg et al., 2016; Unthan et al., 2014). They were inoculated to an initial OD_600_ of 0.1 and incubated overnight at 30 °C at a shaking frequency of 300 rpm and a shaking diameter of 25 mm. The main cultures were inoculated from the precultures after being washed with NaCl solution (0.9% w/v) and resulted in initial OD_600_ of 0.5 for the main culture.

**Figure 1:**
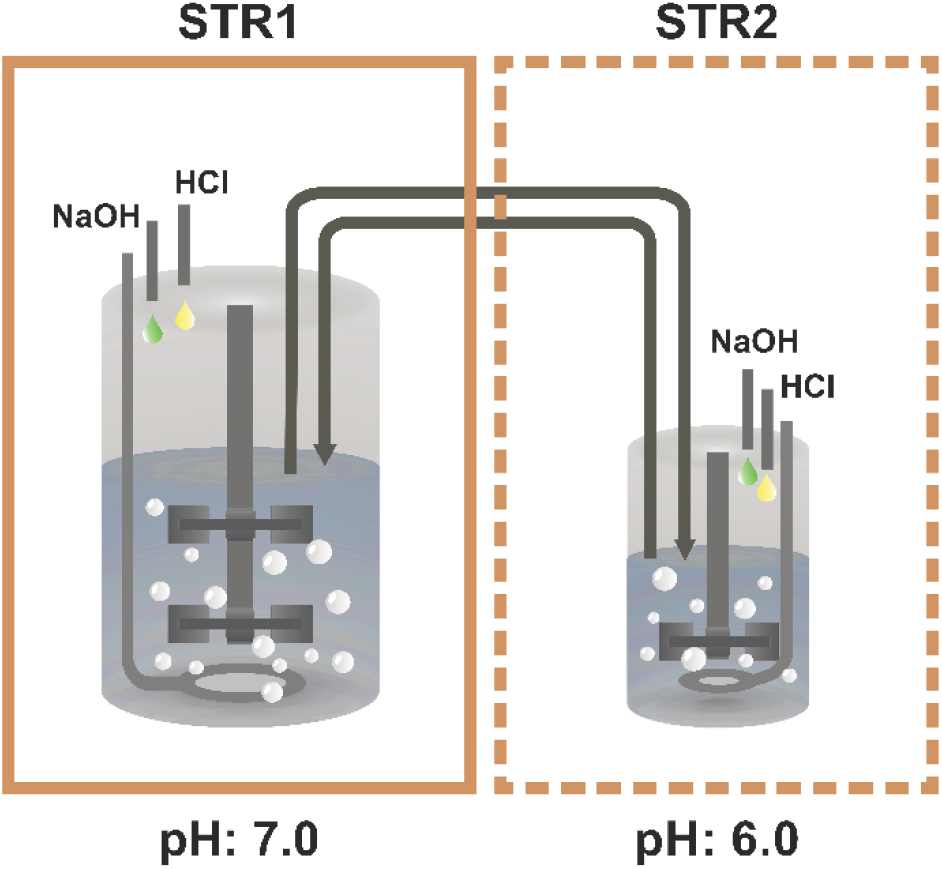
Schematic overview of the two-STR setup. STR1 represents the bulk zone with a pH value of 7, and STR2 represents the disturbance zone with a pH value of 6.

### 2.3 dMSCC – design and principle

The microfluidic system used for dMSCC experiments with *C. glutamicum* described in chapters 3.1 and 3.2 was recently reported by Täuber et al. (Täuber, Golze, et al., 2020). This microfluidic system allows a rapid medium change between two different media. The microfluidic system consists of two inlets between which 14 arrays of cultivation chambers are embedded. The 14 arrays of cultivation chambers are divided into seven different regions, which means that there are always blocks of two cultivation chamber arrays in an oscillation zone. These zones are separated by a channel width of 400 µm. The supply channels have a width of 100 µm and a height of 10 µm. The open-box cultivation chambers are 80 µm x 90 µm x 750 nm in size. For a detailed description of the chip design, the reader is referred to Täuber et al. (Täuber, Blöbaum, Wendisch, & Grünberger, 2021).

A new microfluidic dMSCC design based on the setup described above has been developed for the cultivation of cells under environmental conditions of large-scale bioprocesses (see chapter 3.3). The dMSCC chip in this study contains two parallel, separate cultivation units (Fig. 2A). Each cultivation unit has three inlets for the different medium conditions. Between these inlets are 12 arrays of monolayer cultivation chambers (Fig. 2B). This allows the cultivation of microcolonies up to several hundred cells. The cultivation chamber arrays contained approximately 30 cultivation chambers in one array. The cultivation chambers (Fig. 2C) were designed as open-box monolayer growth chambers (80 µm x 90 µm x 700 nm) to ensure rapid medium exchange via diffusion (Probst et al., 2015). During the dMSCC experiments, the medium can be dynamically switched among three reservoirs, resulting in three distinct cultivation zones. The three zones are separated by a cultivation array-free zone with a width of 400 µm. The zones on the left and right sides of the microfluidic chip serve as control zones, with the central zone of the chip serving as the switching zone. The flow profiles are adjusted to establish the three different flow patterns I – III (Fig. 2D).

**Figure 2:**
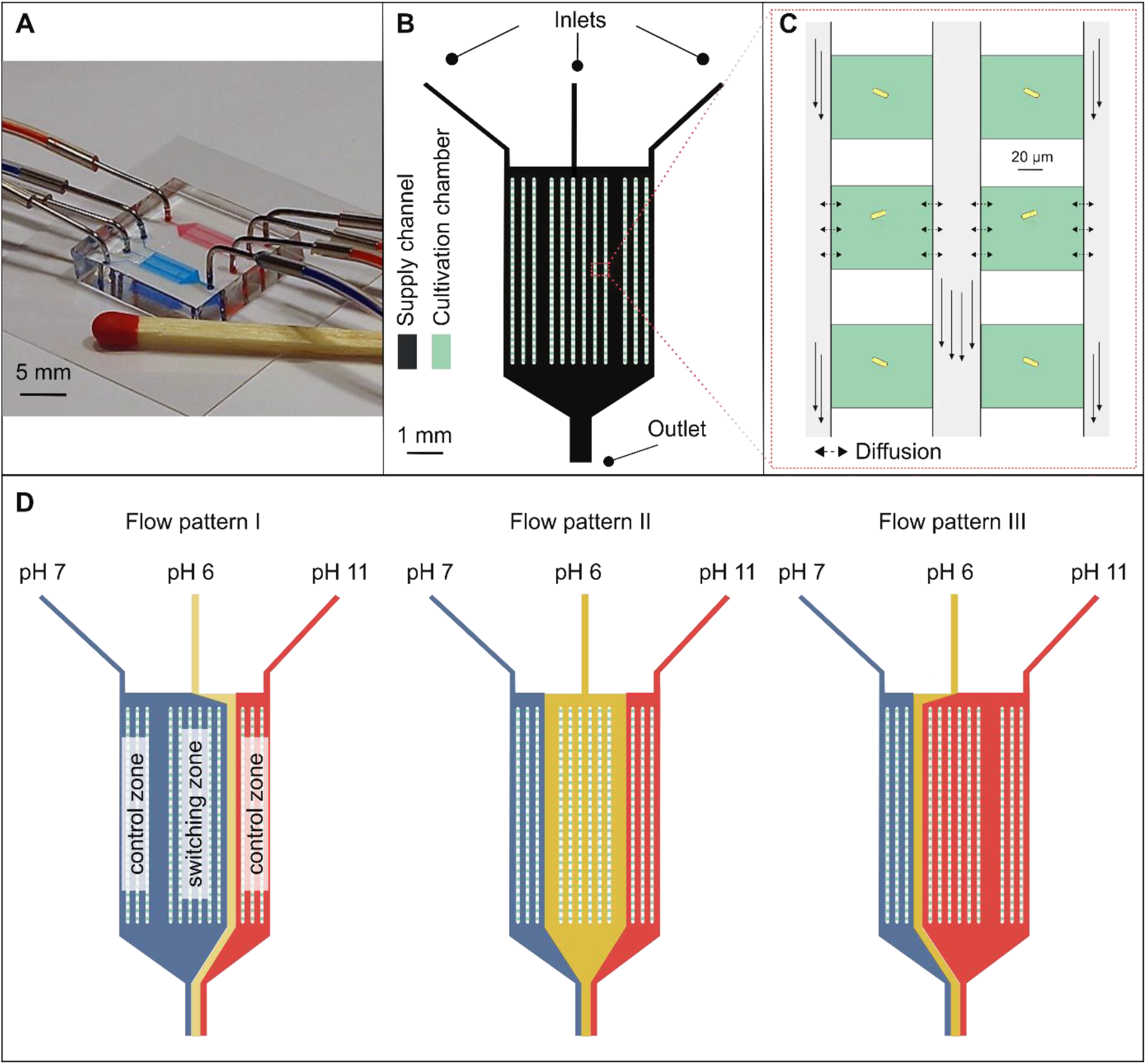
Design of the microfluidic chips for the dynamic cultivation of single cells and microcolonies under three environmental conditions. A) dMSCC chip with three inlets and one outlet per cultivation unit. B) Illustration of the dMSCC design with parallel arrays of cultivation chambers. The three zones are separated by a cultivation array-free zone with a width of 400 µm. C) Monolayer growth chambers. D) Flow pattern at different cultivation conditions in dMSCC.

### 2.4 Microfluidic chip fabrication

Soft lithography was performed for the fabrication of microfluidic chip systems according to the protocol described in Täuber et al. (Täuber, Golze, et al., 2020). In short, the master wafer was covered with PDMS at a ratio of 10:1 between the base and curing agent (Sylgard 184 Silicone Elastomer, Dow Corning Corporation, USA). Afterward, the master wafer was degassed and baked for 2 h at 80 °C. The microfluidic chip was cut off from the master, cleaned and bonded onto a glass slide after 24 s of O_2_ plasma oxidation (Femto Plasma Cleaner, Diener Electronics, Ebhausen, Germany).

### 2.5 Cell seeding procedure in the dMSCC

Cell seeding was performed with a main culture at an OD_600_ of approximately 0.2. The cell suspension was loaded manually with a syringe through the outlet in the microfluidic chip until a sufficient number of cultivation chambers were filled with 1 to 4 cells. The flow of the cell suspension was stopped, the cultivation medium was connected to the microfluidic chip using precision pressure pumps (Fluigent, Jena, Germany), and a pressure of 195 and 40 mbar was set to initiate medium flow.

### 2.6 Oscillation and lifeline experiments

To mimic the oscillation and environmental conditions of large-scale bioprocess experiments (see chapters 3.2 and 3.3), different oscillation parameters were varied for a systematic analysis. These parameters included the oscillation duration *w*, the regeneration time *f* (which is the regeneration time with pH 7), pH oscillation amplitude *A*, total interval duration *T = w+f* and relative oscillation ratio *w*_*rel*_ *= w/T* (Figure 4A, 5A and 6A).

**Figure 3:**
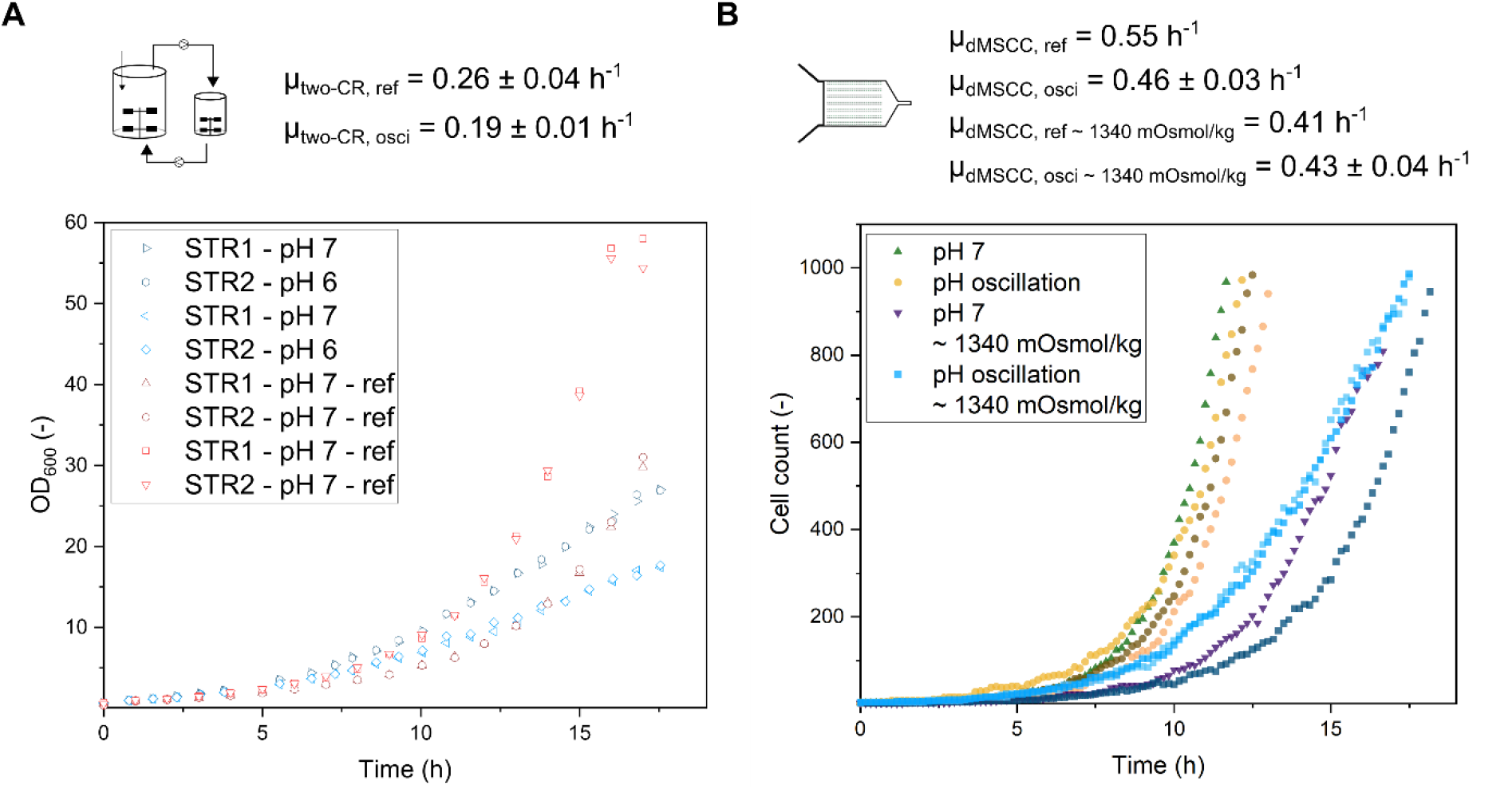
Illustration of the growth curves of *C. glutamicum* from two-CR cultivation and dMSCC for a specific pH oscillation between pH 6 (2.22 min) and pH 7 (7.95 min). A) Displayed is the OD_600_ over the cultivation time for each compartment of the pH oscillation and the control measurement (oscillation between pH 7 and pH 7). DO was controlled at 30 %. B) Displayed is the cell count over the cultivation time for two osmolarities during pH oscillation and at constant pH 7 conditions. Oxygen in the dMSCC system was always available in excess.

**Figure 4:**
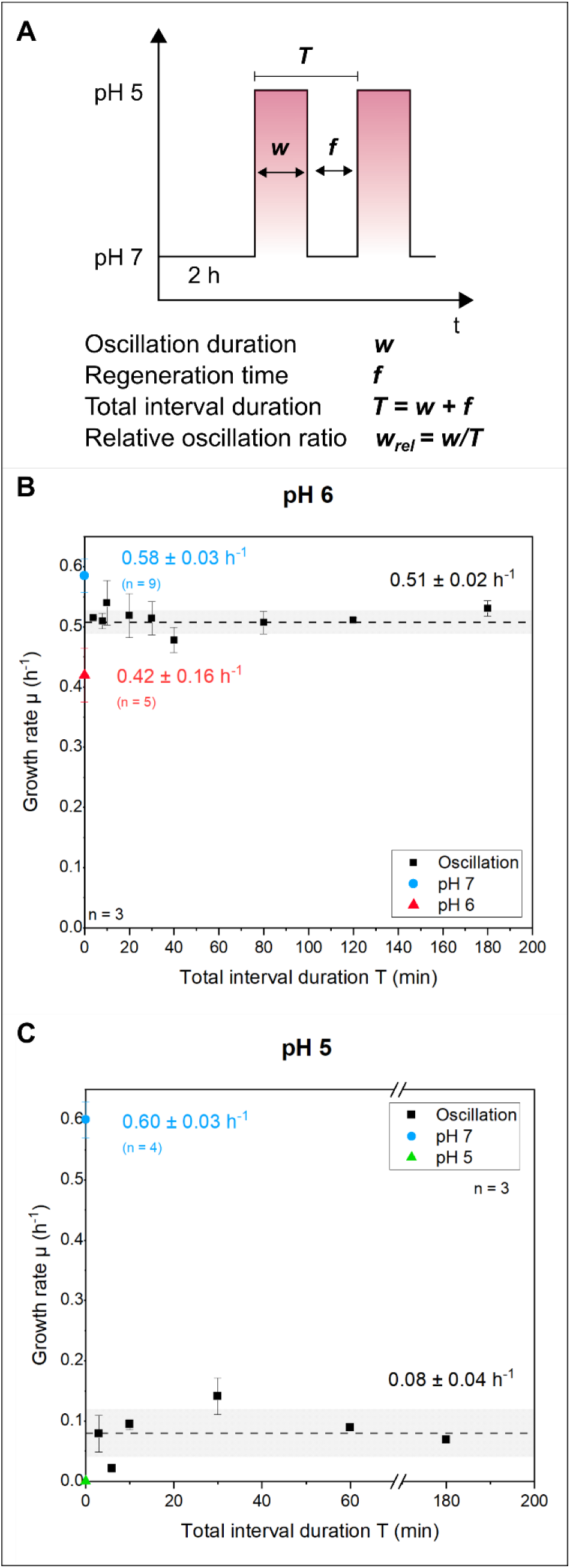
Systematic pH oscillation of *C. glutamicum* with equal oscillation duration *w* and regeneration time *f*. A) Overview of different oscillation parameters. Here, the oscillation duration *w* was equal to the regeneration time *f* so that a relative oscillation ratio *w*_*rel*_ of 0.5 was analyzed. B) The effects of the total interval duration *T* on the growth rate under pH 6/7 oscillation are shown. For control measurements, the growth rate at pH 6 is given in red, and the growth rate at pH 7 is given in blue. C) The effects of the total interval duration *T* on the growth rate at pH 5/7 oscillation are shown. For control measurements, the growth rate at pH 5 is given in green, and the growth rate at pH 7 is given in blue. The standard deviation between the measured growth rates is shown as a gray bar. For each condition, three cultivation chambers were analyzed.

#### 2.6.1 pH oscillation in dMSCC and two-CR scale-down bioreactor

For the dMSCC experiment, we extracted the residence times based on the two-CR cultivation settings (see Table 1) and emulated these conditions. The mean residence time of the cells in the bioreactor was calculated via the pumping rate, which is dependent on the rotation speed and gassing in the large STR. Oscillation parameters were set to an average residence time τ of 2.22 min at pH 6 and 7.95 min at pH 7.

**Table 1:** Mean residence times during the scale-down cultivations.

| Bioreactor |  | Small compartment – STR2 | Large compartment – STR1 |
| --- | --- | --- | --- |
| 1 | Mean residence time $\tau$ – begin | 2.07 min | 7.3 min |
| 1 | Mean residence time $\tau$ – end | 2.09 min | 7.5 min |
| 2 | Mean residence time $\tau$ – begin | 2.33 min | 8.4 min |
| 2 | Mean residence time $\tau$ – end | 2.38 min | 8.6 min |
| | Average mean residence time<br>$T$ | 2.22 min | 7.95 min |

#### 2.6.2 Systematic pH oscillation in dMSCC

For systematic pH oscillation, symmetric oscillations between pH 6 and pH 7 as well as between pH 5 and pH 7 were performed with different total interval durations *T* (3–180 min) and constant relative oscillation ratios *w*_*rel*_ *= 0.5*. Afterward, asymmetric pH oscillations (pH 5 and 7) were performed with varying *w*_*rel*_ from 0.1 to 0.7 and different total interval durations *T* from 10 to 180 min. Finally, pH oscillations between pH 7 and different pH amplitudes *A* (4–6 and 8–11) were performed. Here, the total oscillation duration *T* was set to 20 min, in which the oscillation duration *w* and regeneration phase *f* varied. Different relative oscillation ratios *w*_*rel*_ between 0 and 1 were tested for each oscillation pH value *A*.

#### 2.6.3 Mimicking fluctuating environmental conditions within large-scale bioprocesses

In the final set of experiments, a first fluctuating environmental conditions from large-scale bioprocesses were mimicked by dMSCC. For this purpose, defined intervals of pH 6, 7 and 11 were varied to mimic the gradients of bioreactors that occur through the process of pH control. While concentration profiles for glucose have been modelled by CFD (Haringa, Mudde, & Noorman, 2018), the prediction of pH gradients/profiles in bioreactors has not yet been performed systematically by CFD (Spann et al., 2019). Therefore, a first approximation for potential pH profiles within large-scale bioreactors was constructed based on different pH and concentration profiles related observations and factors extracted from the literature. First, the microenvironment of each cell in bioreactors can switch among different environmental conditions within seconds due to the cell traveling through the bioreactor (Haringa et al., 2018). Second, large-scale bioreactors can require mixing times of up to 2 minutes or more to achieve 95% homogeneity (Junker, 2004; Vrábel, van der Lans, Luyben, Boon, & Nienow, 2000). Third, to simulate the production of acidic compounds or byproduct formation during oxygen limitation, acidification can be applied (Limberg et al., 2017; Zimmermann, Anderlei, Büchs, & Binder, 2006). Fourth, pH control was performed with a pH of 11 to obtain a difference of five pH units between the pH control and the acidified pH value (Reuss et al., 1994). Last, for comparability to the systematic study (Figure 5B), the interval duration was set to 20 minutes. After one oscillation loop, the pH oscillation was repeated until the end of the cultivation. The fluctuating pH profile started with an adaptation phase of 2 hours, followed by the constructed pH oscillation profile shown in Figure 5A. pH oscillation was performed with the parameters shown in Table 2.

**Table 2:**
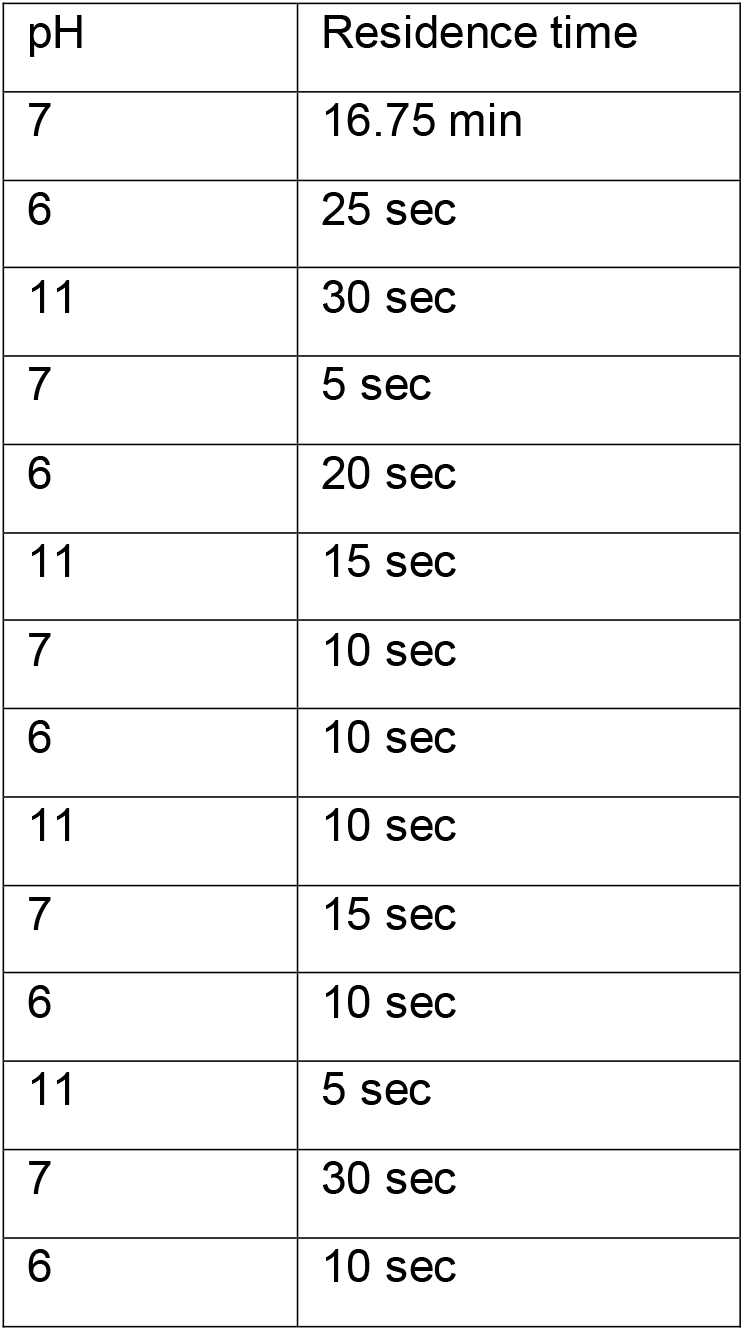
Mimicking fluctuating environmental conditions from the large-scale bioprocess profile. The interval duration of the whole oscillation was set to 20 min. After one loop, the pH oscillation was repeated until the end of the cultivation.

| pH | Residence time |
| --- | --- |
| 7 | 16.75 min |
| 6 | 25 sec |
| 11 | 30 sec |
| 7 | 5 sec |
| 6 | 20 sec |
| 11 | 15 sec |
| 7 | 10 sec |
| 6 | 10 sec |
| 11 | 10 sec |
| 7 | 15 sec |
| 6 | 10 sec |
| 11 | 5 sec |
| 7 | 30 sec |
| 6 | 10 sec |

**Figure 5:**
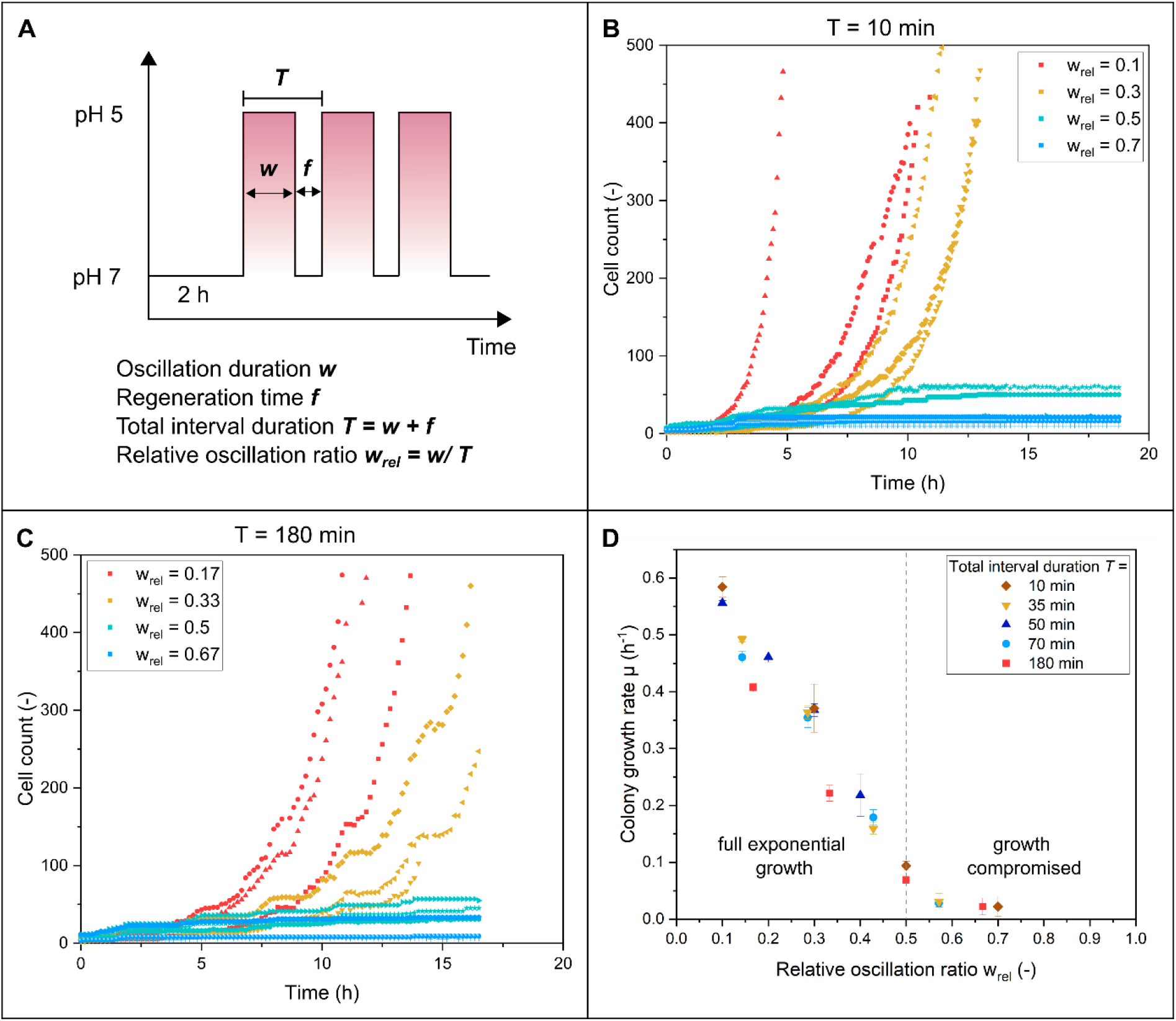
A) Oscillation profile for systematic pH oscillations with two pH values with varying *w*_*rel*_ and *T*. B) Growth curves of *C. glutamicum* with a total interval duration of *T = 10 min* and different relative oscillation ratios w_rel_. Each w_rel_ condition shows three colonies in the same color. C) Growth curves of *C. glutamicum* with a total interval duration of *T = 180 min* and different relative oscillation ratios w_rel_. Each w_rel_ condition shows three colonies in the same color. D) Effect of varying the relative oscillation ratio *w*_*rel*_ for pH 5 oscillations on the growth rate for different interval durations *T*. For *w*_*rel*_ > 0.5, very slow growth was observed, resulting in a maximum of 50 divisions for a cultivation time of 20 h for each colony. Growth curves are shown in Figures S5 – S9 for each total interval duration *T*, and the relative oscillation ratio *w*_*rel*_. For each condition, three cultivation chambers were analyzed.

### 2.7 dMSCC - experimental setup

Live-cell imaging was conducted using a Nikon inverted automated microscope (Nikon Eclipse Ti2, Nikon, Germany). The microscope stage was located in a cage incubator for optimal temperature control (Cage incubator, OKO Touch, Okolab S.R.L., Italy). Inside the cage incubator, the microfluidic device was positioned in an in-house fabricated chip holder. In addition, the setup was provided with a 100× oil objective (CFI P-Apo DM Lambda 100× Oil, Nikon GmbH, Germany), a DS-Qi2 camera (Nikon camera DSQi2, Nikon GmbH, Germany) and an automated focus system (Nikon PFS, Nikon GmbH, Germany) to compensate for thermal drift during long-term microscopy. Eighty cultivation chambers were manually chosen for each experiment and were managed with NIS-Elements Imaging Software (Nikon NIS Elements AR software package, Nikon GmbH, Germany). Pressure-driven pumps (LineUp, Fluigent, Jena, Germany) were used for the generation of different flow profiles. The flow profiles were generated with a specifically tailored cultivation profile implemented in a pump-specific software tool (microfluidic automation tool (MAT), Fluigent, Jena, Germany).

### 2.8 Data analysis of image stacks

The data analysis was performed with the function “analyze particle” from the open-source software Fiji (Schindelin et al., 2012). For this purpose, the phase contrast image sequence was opened, and the cells were separated from the background for each time frame with k-mean clustering. The cluster with the cells was kept, and the background and the intermediate area between the cells was removed and converted to binary images. Cells that were too close together could not be separated by watershed transformation. The integrated function “analyze particle” was then used to calculate the cell number and cell area for each time point. The growth rate was determined graphically by determining the slope of the linear regression from the resulting semilogarithmic plot from the cell count over the cultivation time using OriginPro 2019b (OriginLab Corporation, Northampton, USA). Here, the growth rate was determined from a growing colony inside the cultivation chamber.

## 3. Results and Discussion

### 3.1 A comparison of scales: pH oscillation in two-CR and dMSCC

In the first set of experiments, the growth behavior of *C. glutamicum* was investigated in a two-CR scale-down system under pH oscillation in CGXII medium. STR1 contained a volume of 800 mL, whereas STR2 contained 20% of the total volume (200 mL in STR2). An initial working volume of 1 L was used for both STRs, which oscillated between the two STR compartments. The pH oscillation was started after approximately 7.5 h, as soon as exponential growth was observed, for which the control setpoint was changed to pH 6 in STR2, which was reached after a few minutes only (Fig. S1). The mean residence time of the cells in both compartments was calculated via the pump rate. Based on the calculation, the mean residence time was 7.95 min in STR1 (pH 7) and 2.22 min in STR2 (pH 6). For the pH oscillation in the two-CR system, a growth rate of µ_two-CR, osci_ = 0.19 ± 0.01 h^-1^ was observed in both compartments (Fig. 3A). As a reference, a series of experiments with pH 7 in both compartments was performed. For this purpose, an analogous setup to that used for pH 6/7 oscillation was used. In this experiment, the biomass was also pumped through the two-CRs so that there was biomass exchange between the compartments. Here, STR1 was set to pH 7 and STR2 to pH 7. Cultivation was performed with a cultivation time comparable to the pH 6/7 oscillation. The pH was controlled with HCl or NaOH through a controller, and the DO was controlled at 30% (Fig. S2). A growth rate of µ_two-CR, ref_ = 0.26 ± 0.04 h^-1^ was obtained (Fig. 3A). The lower growth rate for these reference bioreactor experiments compared to the literature values obtained from batch cultivations could be a consequence of the experimental design. Throughout the oscillation phase, the accumulation of sodium chloride salt increased due to pH down and up regulation in the experimental setup. To limit the maximal amount of sodium chloride formed, the total cultivation time was limited to 17 hours of batch cultivation. To ensure sufficient observation time during the exponential batch phase with pH oscillation after the lag-phase, the inoculation density was lowered to an optical density of 0.5. At lower inoculation densities CO_2_ availability could be limiting in the lag-phase. This could have affected the maximum growth rate in the two-CR control experiment. Nevertheless, the growth rate in the pH oscillation experiments in the two-CR system showed a significantly reduced growth rate (−27%) compared to the two-CR reference experiment. This suggests that the effect of the pH oscillation in the two-CR setup is responsible for the reduction.

In the next step, dMSCC was used to determine the growth behavior of *C. glutamicum* under comparable oscillation profiles and aerobic cultivation conditions at 30 °C in CGXII medium. A two-inlet chip (Täuber et al., 2021) was used for the cultivation of cells at two distinct pH values, here pH 6 and 7. For pH oscillation, mean residence times of 2.22 min at pH 6 and 7.95 min at pH 7 were chosen, which resulted from the operation conditions in the two-CR system (see chapter 2.6.1). Using time-lapse microscopy, the growth behavior during dMSCC was recorded for different cultivation chambers. A growth rate of µ_dMSCC, osci_ = 0.46 ± 0.03 h^-1^ was obtained (Fig. 3B). Control experiments, where cell growth was determined with continuous perfusion of CGXII medium at pH 7 resulted in a growth rate of µ_dMSCC,ref_ = 0.55 ± 0 04 h^-1^ for pH 7. The growth of the oscillation experiment in the dMSCC system showed an ∼10% reduction compared to the control experiment at constant pH 7 cultivation conditions.

During two-CR cultivations, medium osmolality increased from ∼700 mOsmol/kg, which represents the starting conditions of the cultivation, to ∼1330 mOsmol/kg at the end of cultivation. A strong increase in osmolarity is a direct consequence of the continuous up and down regulation of the pH value in the two-CRs. These changes were not represented in the dMSCC experiment shown above. Therefore, a set of additional dMSCC experiments was performed in which the medium was adjusted to an osmolarity of ∼1340 mOsmol/kg using NaCl to account for the changes in osmolarity during two-CR cultivations. During the dMSCC experiment under these conditions, a growth rate of µ_dMSCC, osci ∼ 1340 mOsmol/kg_ = 0.32 ± 0.04 h^-1^ was observed for the pH oscillation, and µ_dMSCC, ref ∼ 1340 mOsmol/kg_ = 0.41 ± 0.01 h^-1^ was obtained for the control measurement at constant pH 7 cultivation conditions. The growth of the dMSCC experiment under the combined effect of osmolality and pH oscillations showed a reduction of ∼21% compared to the control measurement.

The obtained growth rates of the reference cultivations in both cultivation systems (two-CR (µ_two-CR, ref_ = 0.26 ± 0.04 h^-1^) and dMSCC (µ_dMSCC, ref_ = 0.55 ± 0.04 h^-1^)) differed most likely due to differences in the setup and cultivation conditions. The obtained growth rates in the dMSCC system are comparable to the growth rates from a constant perfusion microfluidic system (Grünberger et al., 2013; Täuber et al., 2021). The growth rate in the two-CR system is lower than that described in the literature for typical batch cultivations performed in flasks or bench top bioreactors (Graf et al., 2018; Unthan et al., 2014), which may be due to the influence of the shear rates and mechanical burden originating from continuous pumping of the cells between both compartments. Furthermore, this can be explained by the additional current disturbances of the pumping tubes in the reactor and stirring, which discharge CO_2_ from the medium, which has been reported to negatively impact growth, especially at low cell densities (Bäumchen et al., 2007; Blombach, Buchholz, Busche, Kalinowski, & Takors, 2013; Krüger et al., 2019). Furthermore, the “interrupted” oxygen supply in the tube distance between the reactors could affect the growth rate. In addition, in the dMSCC system, there was always an excess of oxygen available due to the gas permeability of PDMS, while the oxygen content in the two-CR was constant at the beginning of cultivation until metabolic oxygen consumption due to population growth exceeded the amount of oxygen available. Thereafter, the oxygen level was kept constant at 30% DO using PID control. The medium availability differed between systems: in the two-CR system, it decreased over the course of cultivation, while in the dMSCC chip, there was always a nutrient excess.

### 3.2 Systematic pH oscillation: growth analysis of C. glutamicum in dMSCC

The dMSCC system offers a wide range of experimental possibilities to perform experiments at different oscillation scenarios, which is impossible in conventional two-CRs. One example is the investigation of the influence of periodic pH fluctuations/oscillations at minutes to seconds intervals on the growth behavior of cells. To demonstrate this, the influence of symmetric pH oscillation on the growth rate of *C. glutamicum* was further investigated (Fig. 4). The aim of these experiments was to investigate the effects of different interval durations *T* on the growth of *C. glutamicum* during pH 6/7 and 5/7 oscillations when exposed to equal ratios of the oscillation duration *w* and regeneration time *f* (Fig. 4A). Figure 3B shows the growth rate as a function of the total interval duration *T*. A growth rate of µ = 0.51 ± 0.02 h^-1^ was observed for pH 6/7 oscillations of different total interval durations *T* between 4 and 180 min (Fig. 4B & S3). The total interval duration *T* had no significant influence on the growth rate during the oscillation. During symmetric oscillations at pH 5/7, cellular growth was significantly affected, resulting in compromised growth throughout the cultivation. For almost all colonies of all oscillation intervals, cell growth ended after a few division events (Fig. S4 – green and blue curves). One colony at an oscillation duration of *w = 3 min* (*T = 6 min*) was observed to grow slowly but exponentially over the course of cultivation (Fig. S4 – red curve). For consistency, growth rates were estimated by exponential fitting in analogy to the data obtain in Figure 4B and resulted in µ = 0.08 ± 0.04 h^-1^ (Fig. 4C). The reference cultivation at pH 7 yielded µ = 0.58 ± 0.03 h^-1^, which was consistent with the other reference cultivation at this pH (Täuber et al., 2021; Unthan et al., 2014). The cultivations at pH 5 showed no cell division, indicating that *C. glutamicum* could not grow at this pH, which is in agreement with previously reported cultivations at this pH (Täuber et al., 2021).

After observing the effect of the total interval duration *T* at a constant oscillation ratio (*w*_*rel*_ *= 0.5*), the following series of experiments investigated the variation in the oscillation ratio *w*_*rel*_ at pH 5 with a combination of different total interval durations *T* (Figure 5A). A stronger influence on growth performance was observed compared to that of pH 6 based on previous measurements (see chapter 3.1). Oscillation ratios in bioprocesses are highly variable due to the movement of cells through the zones of the bioreactor. These zones influence the growth behavior of individual cells. Two key questions were as follows: How does the oscillation rate *w*_*rel*_ affect the growth rate *µ* and do different oscillation ratios *w*_*rel*_ affect the influence of the total interval time *T* on the growth rate? Figure 5B and C show the growth curves from total interval durations *T = 10 min* and *T = 180* min with different relative oscillation ratios *w*_*rel*_. The growth decreased as the relative oscillation ratio *w*_*rel*_ increased. Plateaus appear in the growth curves at long total interval durations, such as *T = 180 min*. However, these lag phases are longer than the pH stress value, so no clear correlation between pH oscillation and lag phase have been observed. This phenomenon will be further investigated in the future. Figure 5D shows the correlation between the growth rate and relative oscillation ratio *w*_*rel*_ for different total interval durations *T* between 10 and 180 min. The growth rate correlated reciprocally with the relative oscillation ratio *w*_*rel*_ and decreased with increasing relative oscillation ratios *w*_*rel*_. No correlation was found for relative oscillation ratios *w*_*rel*_ greater than 0.5. At high relative oscillation ratios, not all daughter cells continued to grow, so no exponential growth was observed. Here, only a few division events (< 20) were observed over a cultivation time of 20 h. One could speculate that longer oscillation durations *w* at pH 5 lead to an imbalanced intracellular pH and problems in maintaining pH homeostasis (Fig. 5B & 5C – blue and green curves). Strikingly, the total interval duration *T* seems to have no important impact. In addition, the decrease in growth rate was examined in more detail to determine if it was directly related to the growth (pH 7) and nongrowth (pH 5) phases. For this purpose, the experimental data shown in Fig. 5D were compared with theoretically calculated data. The theoretical data consider only the effect of growth at pH 7 in the regeneration times *f* and nongrowth at pH 5 in the oscillation durations *w*. These data were obtained using the equation in Fig. S10. A growth rate of µ_pH 5_ = 0 h^-1^ at pH 5 and of µ_pH 7_ = 0.58 ± 0.03 h^-1^ at pH 7 was assumed. When comparing the theoretical data with the experimental data, it is clear that the experimental data show a greater decrease in the growth rate. It can be concluded that the experimental data obtained here cannot be explained by the relationship between the growth phase and the nongrowth phase alone. Therefore, it can be suspected that the cells underwent a short pH 5-induced lag phase immediately following the pH 5 nongrowth phase. The formation of pH-induced lag phases was already observed after single pH stress pulses between 2 and 9 h (Täuber et al., 2021). The intracellular adaptation to the oscillation may cost time and metabolic energy, which leads to a reduction in the growth rate.

In the next set of experiments, the growth behavior of *C. glutamicum* was investigated under different oscillating pH conditions. Oscillation between pH 7 and oscillation amplitudes *A* of pH 4–6 and 8–11 was performed with varying relative oscillation ratios *w*_*rel*_ (difference between oscillation duration *w* and regeneration time *f* (Fig. 6A)) and a constant total interval duration (*T = 20 min*). Figure 5B shows the growth rate for each oscillation as a function of the relative oscillation ratio *w*_*rel*_. The growth rate decreased significantly with increasing relative oscillation ratio *w*_*rel*_ for all pH oscillation amplitudes *A* used. The growth rate correlated reciprocally with the relative oscillation ratio *w*_*rel*_ for all pH amplitudes *A* shown (Fig. 6B). At pH 6, the growth rate decreased with increasing relative oscillation ratios w_rel_ down to a growth rate of µ = 0.4 h^-1^. A similar effect was observed for the other pH oscillation amplitudes *A*. Here, the growth rates dropped to 0 h^-1^ at a defined relative oscillation ratio *w*_*rel*_, e.g., at pH 9, at a relative oscillation *w*_*rel*_ of 0.8 or higher, the growth rate was lower than µ = 0.1 h^-1^ and decreased to 0 h^-1^ as the relative oscillation ratio *w*_*rel*_ increased. Here, the cells showed growth in area but did not necessarily divide anymore during the timeframe of observation. The growth rates obtained from dMSCC cultivation at pH 4 and 11 showed a rapid decrease; therefore, this experiment was analyzed only for relative oscillation ratios *w*_*rel*_ in the range of 0 and 0.125.

**Figure 6:**
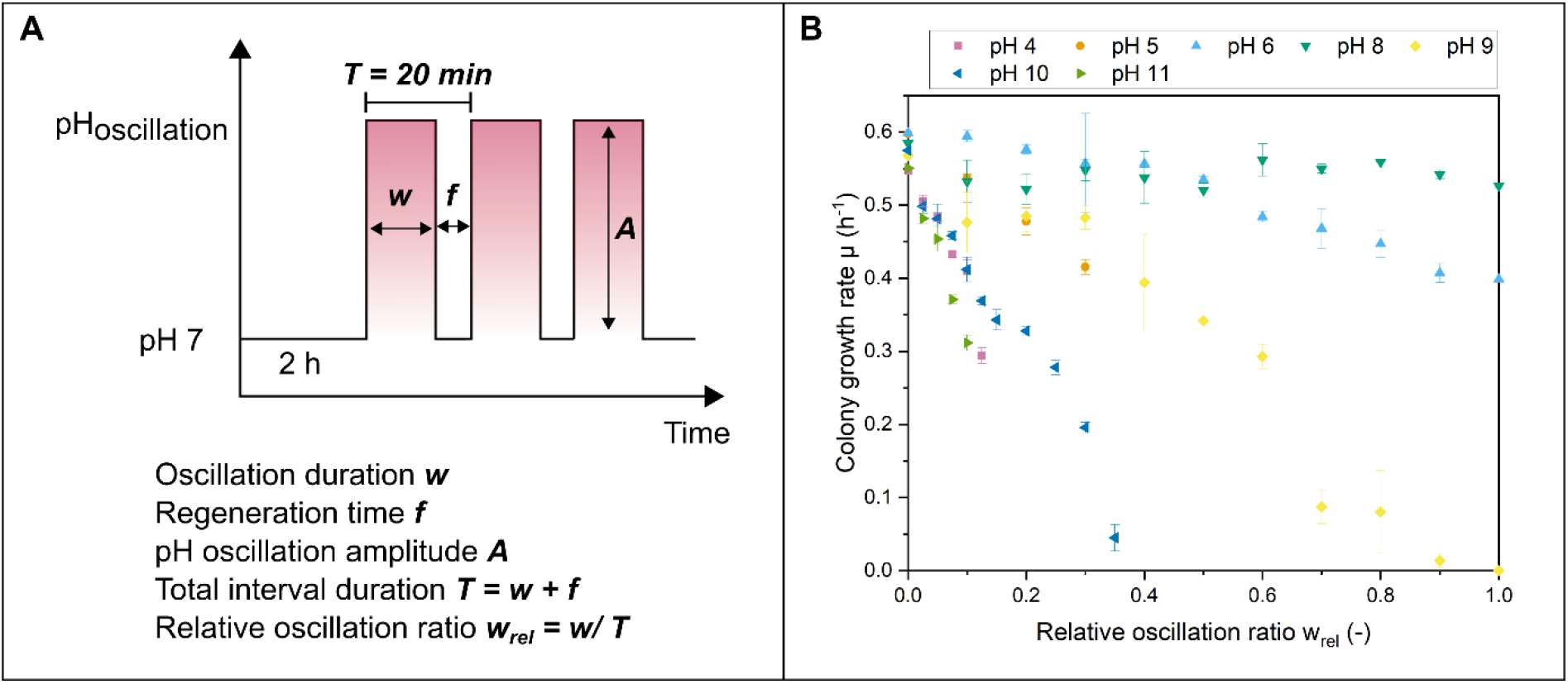
A) Oscillation profile for systematic pH oscillations with two pH values where the total interval duration *T* was set to 20 min. B) Growth rate depended on the different relative oscillation ratios *w*_*rel*_. The relative oscillation ratio *w*_*rel*_ was applied with a total interval duration *T* of 20 min for pH stress amplitudes *A* 4-11. For each condition, three cultivation chambers were analyzed.

A linear fit was applied to each pH amplitude *A* as a qualitative comparison (Fig. S11). The oscillation indicator, the slope, differed between the tested pH amplitudes. The stress level can thus be ordered as follows: pH 8 < pH 6 < pH 9 < pH 5 < pH 10 < pH 4 < pH 11. These oscillations triggered a lag phase (Fig. S3 – blue and orange curve and Fig. S9 red and orange curves), which can be explained by the intracellular adaptation to the oscillation, which requires time and metabolic energy (Täuber et al., 2021). The lag phase was observed at both high and low oscillation frequencies. The internal pH of a cell is maintained by pH homeostasis, which involves different constitutive and regulatory mechanisms to overcome pH stress. In *C. glutamicum*, the internal pH is 7.5 ± 0.5 at environmental pH values between 5.5 and 9. Beyond this range, the internal pH collapses, and pH homeostasis fails. Under acidic conditions, mycothiol, which protects cysteine and methionine residues of proteins from ROS by reversible S-mycothiolations, could be affected by pH oscillation, and fast protein regeneration would not be possible (Liu et al., 2016). Acidic stress may require biochemical reaction mechanisms (e.g., mycothiol, protein refolding, iron transport, etc.) to respond to the stress pulse. Under alkaline conditions, ROS protection is not necessary, as ROS do not accumulate at significant concentrations (Follmann et al., 2009). Here, it is possible that a genetic response to pH stress is necessary, which would require time. Mrp-type Na^+^/H^+^ antiporters play a dominant role in the resistance of *C. glutamicum* cells to alkaline pH. For example, in the case of osmotic stress (Wood et al., 2001), first, a fast response by transport and uptake of compatible substances from the medium is carried out, and then a genetic response is triggered.

In summary, the dMSCC system allowed a detailed and accurate analysis of the growth behavior of *C. glutamicum* colonies under different oscillation parameters, such as oscillation amplitude *A* and total interval duration *T*. Rapid and targeted oscillations can be performed without affecting other cultivation parameters (such as substrate and oxygen availability) at high temporal resolution, which is not possible in traditional scale-down reactors.

### 3.3 Mimicking fluctuating environmental conditions within large-scale bioprocesses in dMSCC

To demonstrate the potential of dMSCC, we mimicked the fluctuating environmental conditions in large-scale bioreactors created by the process of pH control to study cell growth behavior under mimicked conditions. A three inlet dMSCC system (see chapter 2.3 and Fig. 2) was used for rapid and precise oscillation of the medium between pH 6, 7 and 11 at varying oscillation duration *w*, regeneration time *f* and amplitude *A*. The fluctuating environmental conditions in large-scale bioprocesses were translated into a pH concentration profile to perform dMSCC experiments (Fig. 7A). In short, the profile consisted of different pH and concentration profiles related observations. For example, mixing times in large-scale bioreactors require up to 2 min to achieve 95% heterogeneity in the bioreactor, dependent on the size and geometry of the bioreactor (Haringa et al., 2018), and acidification was indicated as a first pH change when simulating acidic products. The profile was repeated every 20 min. Details on the design of the profile, which mimics the fluctuating environmental conditions in large-scale bioprocesses, can be found in chapter 2.6.3.

**Figure 7:**
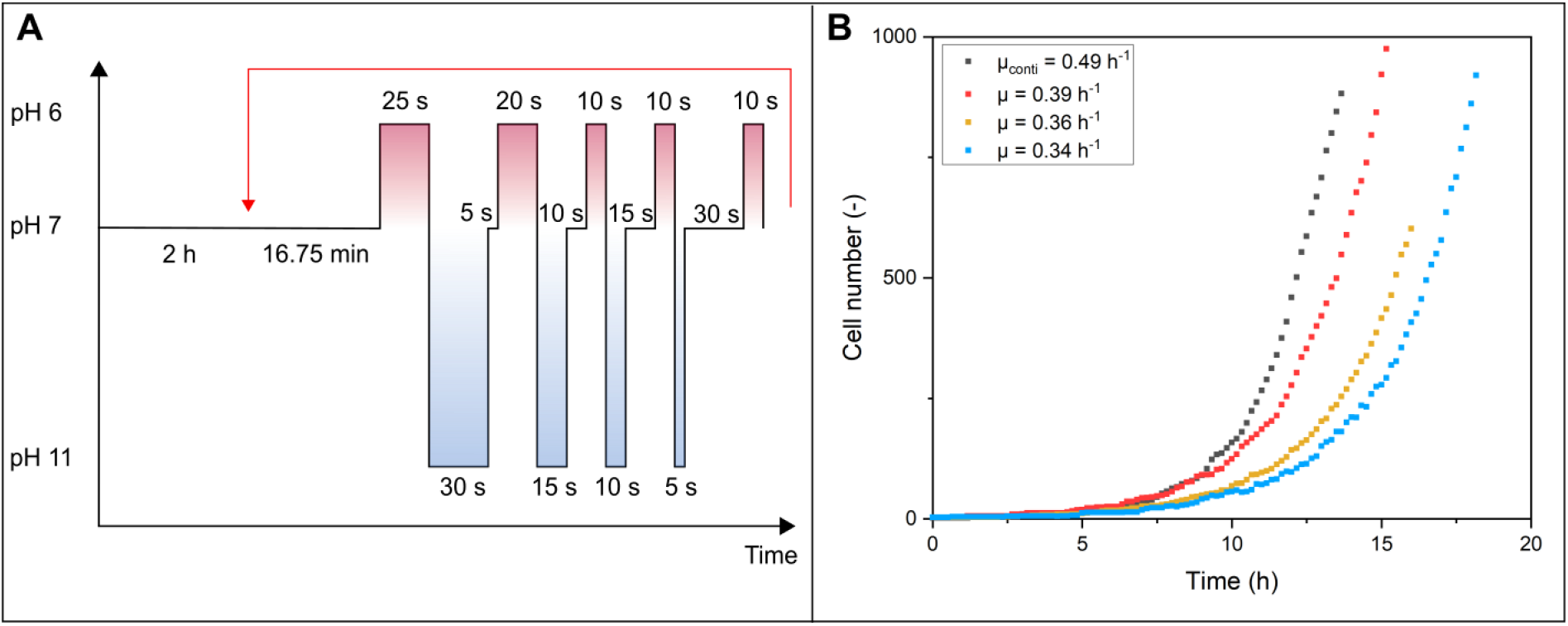
Growth of *C. glutamicum* under a coarse-grained pH lifeline. A) Representative cell lifeline during pH adjustment in a large-scale bioreactor. B) Growth curves of *C. glutamicum* in dMSCC showing the cells’ pH profile of three colonies (red, yellow, blue). The control experiments of cells cultivated at pH = 7 are shown in black.

Exponential growth was observed for all colonies under mimicked fluctuating environmental conditions in large-scale bioprocesses. A growth rate of µ = 0.36 ± 0.02 h^-1^ was found for the cells exposed to the mimicked environmental conditions (Fig. 7B). The control experiments, in which cell growth was determined with continuous perfusion of CGXII medium at pH 7, resulted in a growth rate of µ = 0.49 h^-1^. The growth under fluctuating environmental conditions in the large-scale bioprocess experiment showed an ∼ 27% reduction in growth compared to the constant-perfusion cultivation conditions at pH 7. Based on the results of the growth rate of *C. glutamicum* dependent on the relative oscillation ratios *w*_*rel*_ for different pH values *A* (see Fig. 6B), a growth rate reduction of ∼ 18% would be expected for oscillations with pH 11 and a relative oscillation ratio *w*_*rel*_ of 0.05, which is a comparable relative oscillation ratio *w*_*rel*_ to the mimicked fluctuating environment profile shown here. For the applied mimicked pH profile with a *w*_*rel,pH 11*_ of 0.05, a similar growth rate reduction would be assumed due to the comparable oscillation ratio if the oscillation duration of pH 6 can be neglected, as it has already been shown in chapter 3.2 that for relative oscillation ratios *w*_*rel*_ smaller than 0.2, no effect on the growth rate was observed with pH 6/7 oscillation. This implies that first, cell growth under the mimicked conditions was not only defined by the pH 11 oscillation ratio, and second, the additional pH 6 oscillation ratio in combination with the pH 11 oscillation had a significant negative synergistic effect on the growth rate.

To summarize, the dMSCC system can be used to experimentally mimic concentration profiles from bioreactors to analyze the effects of changing pH conditions on the cell growth rate.

The control measurements of the pH profile and the scale comparison experiments ranged from µ = 0.49 h^-1^ to µ = 0.55 h^-1^. The differences in the control measurements can be explained by the variability of the individual cells, which can show day-to-day variance.

### 3.4 Comparative discussion of two-CR and dMSCC systems

The results and application examples demonstrated that dMSCC offers the opportunity to mimic fluctuating environmental conditions within large-scale bioprocesses. In the following both technologies, the two-CR and dMSCC, will be compared in terms of cultivation setup, environmental control and analytics (Fig. 8).

**Figure 8:**
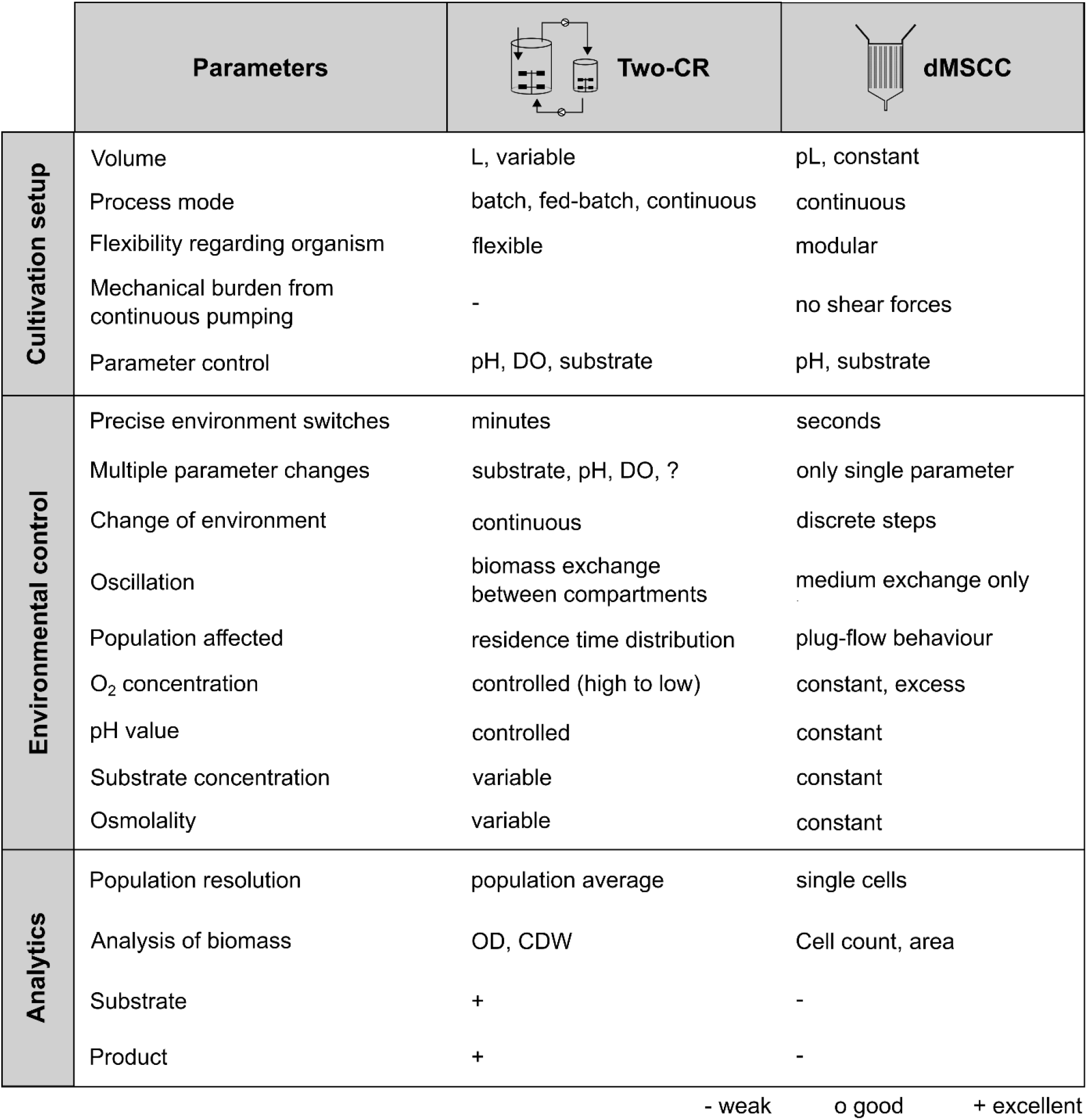
Comparison of the two-CR and dMSCC systems. The aspects were divided into three categories: cultivation setup, environmental control and analytics.

The cultivation setup of the two systems differs in volume (L vs. pL) and process control. Batch, fed-batch and continuous cultivations can be performed in the two-CR system. Due to the flexibility in the configuration of the different compartments, analyses of different parameters, such as biomass and metabolism, can be performed, which allows the cultivation of various organisms and strains. In the two-CR system, shear forces and mechanical burden are present due to continuous pumping between the compartments, which can influence the growth rate and productivity of cells. During cultivation, various process parameters, e.g., pH value and DO, can be controlled. The corresponding devices, e.g., pH meters, determine the response time of the respective probes (time until stable measurement) and the reaction time of the control loop, which should be considered for the oscillations.

Currently, only continuous cultivations can be performed in the dMSCC system. The cultivation chamber geometry of the dMSCC setup has fixed dimensions. Therefore, the setup is restricted for the cultivation of organisms and strains of the same height (Binder et al., 2016; Grünberger et al., 2013). Setups with different chamber heights are necessary for the cultivation of larger cells, such as mammalian cell lines (Schmitz et al., 2021). In the dMSCC system, there are negligible shear forces because the cells are trapped in the cultivation chambers, and a medium supply is obtained only per diffusion (Westerwalbesloh et al., 2015). Control of the pH value and DO concentration is possible by adjusting the medium. Oscillations in the pH value are possible only by changing the feed medium by adjusting the corresponding pH value in the medium.

Additional differences between the two setups can be found within the field of environmental control. In the two-CR setup, oscillations in timeframes between 0.5 and 11 min can be performed (Larsson & Enfors, 1988; Namdev, Thompson, & Gray, 1992). In the two-CR system used here, the shortest possible/documented residence time in STR2 is 0.8 min. The setup allows multiple parameters to be studied simultaneously in one setup. By changing the setup configurations of the scale-down reactors, different parameters can be studied in STR-STR/PFR, from 3 parameter studies, e.g., substrate, DO concentration and pH value (Olughu et al., 2020), to 2 parameter studies, e.g., the gradients of substrates and DO concentration (Lemoine, Maya Martínez-Iturralde, Spann, Neubauer, & Junne, 2015; Limberg et al., 2016), to one parameter study, e.g., substrate in complex medium (Baez, Flores, Bolívar, & Ramírez, 2011; Lemoine et al., 2016). The changes in conditions are steady so that continuous oscillations between two parameters can be performed. Oscillation occurs through the exchange of biomass between the different compartments, exposing only some of the cells to oscillation at a time. This results in a residence time distribution of the affected population. Oscillatory cultivation ends at the latest when a given total volume is reached. DO and pH can be controlled. Specifically, during pH oscillation, there is an increase in osmolarity in the two-CR due to the pH control process.

In the dMSCC system, a more precise environmental condition can be implemented because the medium changes can be realized within a few seconds, allowing medium oscillation times as low as 5 seconds. Currently, oscillations can be investigated with only one environmental parameter (Täuber, Golze, et al., 2020). These oscillations can be performed quickly and precisely in two or three discrete values of a single parameter. During cultivation, the cells are fixed in one place, and the medium is constantly exchanged, exposing all cells to oscillation simultaneously. This affects the population in a similar way to plug flow behavior. The analysis of the cells under oscillating conditions can be performed for several generations depending on the chamber size and thus exceeds the timespan of typical two-CR batch cultivations. Oxygen is always available in excess in the dMSCC system, as the system is gas permeable. Another important parameter during cultivation is osmolarity. In the two-CR system, an increase in osmolarity is due to the pH control process, which must be additionally taken into account in the dMSCC system, where it is constant.

Further differences between the two technologies are in the field of analytics. Two-CR systems operate on a bulk scale, so the heterogeneity of the environment can be studied, not the cell-to-cell heterogeneity within the population (Kovárová-Kovar & Egli, 1998). Therefore, the insights mostly based on averaged population data (Lindström & Andersson-Svahn, 2010; Templer & Ces, 2008). The study of cell-to-cell variations in two-CRs has been demonstrated but not performed systematically. Brognaux et al. combined a two-CRs with flow cytometry to gain deeper insights regarding cell-to-cell heterogeneity within traditional scale-down reactors (Brognaux et al., 2013). In the two-CR system, many parameters can be measured both online, e.g., pH value and DO concentration, and offline, e.g., biomass and metabolism during cultivation. Information on the absolute amount of products, product yield, metabolism, etc. can thus currently only be studied in scale-down bioreactors.

In the dMSCC system, the behavior of individual cells can be studied at high spatiotemporal resolution through live-cell imaging, enabling the investigation of single-cell dynamics. The analysis in the dMSCC system is currently restricted to image-based analysis of cell behavior and thus to the determination of growth rate, morphology and fluorescence coupled metabolic processes (Mustafi et al., 2014; Täuber, Golze, et al., 2020).

The comparison between the two-CR and dMSCC approaches shows that dMSCC has the potential to be used as a microfluidic single-cell scale-down reactor. Single-cell scale-down reactors serve as a complementary technology to CRs regarding their possibilities for scale-down studies, as single-cell scale-down reactors have high potential to be used for high frequency oscillating scale-down studies and for cellular lifeline studies of large-scale bioreactors (Ho et al., 2019). CRs should preferably be used for multiparameter studies, which are not currently possible in the dMSCC system, as already shown by Käß et al. (Käß, Hariskos, et al., 2014). Another field of application may be “omics”-coupled scale-down studies (Delvigne, Takors, Mudde, van Gulik, & Noorman, 2017; Wehrs et al., 2019) due to the broad spectrum of different analytical methods, which can be performed only with the CR system. The dMSCC system can be used for systematic studies of oscillating media parameters to investigate cellular responses with respect to various variable oscillation parameters (Täuber, Golze, et al., 2020), which is not possible for traditional CR systems.

The dMSCC system needs to be further optimized so that the influence of multiple parameters can be studied simultaneously in one device to mimic bioprocesses. The influence of different feast and famine cycles, different medium compositions and concentrations on the cellular behavior at the level of individual microbial cells can be investigated. An important parameter is the DO, which thus far cannot be controlled and measured in the chip. This property must also be able to be varied in the chip to better represent the bioprocess.

Insights into bioprocess heterogeneity during different oscillations could be gained by using single-cell growth channels, often referred to as mother machines (Wang et al., 2010), instead of the cultivation chamber used in this work. In mother machines, single cells can be grown for several generations, and detailed dynamic studies can be performed. This allows quantification of single-cell growth (i.e., elongation and time to cell division) over many generations and makes it possible to determine whether and to what extent oscillations affect partial nonuniformities and production. In future, two-CR and dMSCC can complement each other. Using two-CRs to investigate biomass and productivity upon overall gradients offers the opportunity to gain a deeper understanding of cellular responses to gradient formation. dMSCC can supplement these experiments by providing insights into cell-to-cell heterogeneity regarding fast and defined fluctuating environmental conditions such as medium compounds or pH.

## 4. Conclusion

The study of fluctuating environments and gradients in bioreactors can be very complex and accordingly difficult to analyze. Until now, analysis of the effects of gradients on cellular performance in CR setups has required laboratory-scale experiments. Recently, microfluidic cultivation systems have become available that could complement these methods. In this study, first, a growth comparison of *C. glutamicum* in a two-CR and dMSCC was conducted for a specific pH oscillation between pH 6 and pH 7. Here, a reduction of ∼ 21% in growth rate was observed in the dMSCC system, and a reduction of ∼ 27% was observed in the two-CR system compared to cultivation of *C. glutamicum* at standard conditions (pH = 7). A stronger decrease in the growth rate was observed for the cultivation of cells in the two-CRs, which could be attributed to reactor-specific influences. Furthermore, the dMSCC system was used for a detailed analysis of the effects of different periodic pH 6/7 and 5/7 oscillations at the single-cell level, which is not possible with conventional bioreactors, since fluctuating environments can be implemented more precisely in this system. A systematic pH oscillation study was performed with varying relative oscillation ratios, total interval durations and different pH oscillation amplitudes. The results showed a significant effect of the different pH oscillations on the growth rate of *C. glutamicum*. The experiments were used to demonstrate that the presented microfluidic system can be used in the future as a scale-down tool and to show which information can be obtained by these systems compared to two-CR systems. Two-CR and dMSCC can complement each other. The use of two-CRs to study biomass and productivity upon overall gradients offers the opportunity to gain a deeper understanding of cellular responses to gradient formation. dMSCC can supplement these experiments by providing insights into cell-to-cell heterogeneity regarding fast and defined fluctuating environmental conditions such as medium compounds or pH. The dMSCC system can be used in the future to understand the behavior of strains under defined oscillatory conditions forms the basis for the development of future process control strategies or the engineering of more robust production strains regarding existing gradients in industrial scale bioreactors.

## Acknowledgements

Parts of this work were performed at the cleanroom facilities of the Department of Biophysics and Nanoscience as well as the Department for Physics of Supramolecular Systems and Surfaces at Bielefeld University. The authors are thankful for all the help and support.

## References

Amanullah, A., McFarlane, C. M., Emery, A. N., & Nienow, A. W. (2001). Scale-down model to simulate spatial pH variations in large-scale bioreactors. Biotechnology and Bioengineering, 73(5), 390–399. https://doi.org/10.1002/bit.1072

Baez, A., Flores, N., Bolívar, F., & Ramírez, O. T. (2011). Simulation of dissolved CO_2_ gradients in a scale-down system: A metabolic and transcriptional study of recombinant Escherichia coli. Biotechnology Journal, 6(8), 959–967. https://doi.org/10.1002/biot.201000407

Bäumchen, C., Knoll, A., Husemann, B., Seletzky, J., Maier, B., Dietrich, C., Amoabediny, G., & Büchs, J. (2007). Effect of elevated dissolved carbon dioxide concentrations on growth of Corynebacterium glutamicum on D-glucose and L-lactate. Journal of Biotechnology, 128(4), 868–874. https://doi.org/10.1016/j.jbiotec.2007.01.001

Becker, J., Kuhl, M., Kohlstedt, M., Starck, S., & Wittmann, C. (2018). Metabolic engineering of Corynebacterium glutamicum for the production of cis, cis-muconic acid from lignin. Microbial Cell Factories, 17(1), 115. https://doi.org/10.1186/s12934-018-0963-2

Becker, J., & Wittmann, C. (2017). Industrial Microorganisms: Corynebacterium glutamicum. In C. Wittmann & J. C. Liao (Eds.), Industrial Biotechnology (Vol. 3, pp. 183–220). Weinheim, Germany: Wiley-VCH Verlag GmbH & Co. KGaA. https://doi.org/10.1002/9783527807796.ch6

Binder, D., Probst, C., Grünberger, A., Hilgers, F., Loeschcke, A., Jaeger, K.-E., Kohlheyer, D., & Drepper, T. (2016). Comparative Single-Cell Analysis of Different E. Coli Expression Systems during Microfluidic Cultivation. PloS One, 11(8), e0160711. https://doi.org/10.1371/journal.pone.0160711

Blombach, B., Buchholz, J., Busche, T., Kalinowski, J., & Takors, R. (2013). Impact of different CO_2_/HCO_3_-levels on metabolism and regulation in Corynebacterium glutamicum. Journal of Biotechnology, 168(4), 331–340. https://doi.org/10.1016/j.jbiotec.2013.10.005

Bothun, G. D., Knutson, B. L., Berberich, J. A., Strobel, H. J., & Nokes, S. E. (2004). Metabolic selectivity and growth of Clostridium thermocellum in continuous culture under elevated hydrostatic pressure. Applied Microbiology and Biotechnology, 65(2), 149–157. https://doi.org/10.1007/s00253-004-1554-1

Brognaux, A., Han, S., Sørensen, S. J., Lebeau, F., Thonart, P., & Delvigne, F. (2013). A low-cost, multiplexable, automated flow cytometry procedure for the characterization of microbial stress dynamics in bioreactors. Microbial Cell Factories, 12, 100. https://doi.org/10.1186/1475-2859-12-100

Champagne, C. P., Gardner, N., & Doyon, G. (1989). Production of Leuconostoc oenos Biomass under pH Control. Physical Review Letters, 55(10), 2488–2492. https://doi.org/10.1128/aem.55.10.2488-2492.1989

Cortés, J. T., Flores, N., Bolívar, F., Lara, A. R., & Ramírez, O. T. (2016). Physiological effects of pH gradients on Escherichia coli during plasmid DNA production. Biotechnology and Bioengineering, 113(3), 598–611. https://doi.org/10.1002/bit.25817

Crater, J. S., & Lievense, J. C. (2018). Scale-up of industrial microbial processes. FEMS Microbiology Letters, 365(13), 42. https://doi.org/10.1093/femsle/fny138

Delvigne, F., Takors, R. [Ralf], Mudde, R., van Gulik, W., & Noorman, H. (2017). Bioprocess scale-up/down as integrative enabling technology: From fluid mechanics to systems biology and beyond. Microbial Biotechnology, 10(5), 1267–1274. https://doi.org/10.1111/1751-7915.12803

Follmann, M., Ochrombel, I., Krämer, R., Trötschel, C., Poetsch, A., Rückert, C., Hüser, Andrea; Persicke, M., Seiferling, D., Kalinowski, J., & Marin, K. (2009). Functional genomics of pH homeostasis in Corynebacterium glutamicum revealed novel links between pH response, oxidative stress, iron homeostasis and methionine synthesis. BMC Genomics, 10, 621. https://doi.org/10.1186/1471-2164-10-621

Graf, M., Zieringer, J., Haas, T., Nieß, A., Blombach, B., & Takors, R. (2018). Physiological Response of Corynebacterium glutamicum to Increasingly Nutrient-Rich Growth Conditions. Frontiers in Microbiology, 9, 2058. https://doi.org/10.3389/fmicb.2018.02058

Grünberger, A., van Ooyen, J., Paczia, N., Rohe, P., Schiendzielorz, G., Eggeling, L., Wiechert, W., Kohlheyer, D., & Noack, S. (2013). Beyond growth rate 0.6: Corynebacterium glutamicum cultivated in highly diluted environments. Biotechnology and Bioengineering, 110(1), 220–228. https://doi.org/10.1002/bit.24616

Hansen, G., Johansen, C. L., Marten, G., Wilmes, J., Jespersen, L., & Arneborg, N. (2016). Influence of extracellular pH on growth, viability, cell size, acidification activity, and intracellular pH of Lactococcus lactis in batch fermentations. Applied Microbiology and Biotechnology, 100(13), 5965–5976. https://doi.org/10.1007/s00253-016-7454-3

Haringa, C., Mudde, R. F., & Noorman, H. J. (2018). From industrial fermentor to CFD-guided downscaling: what have we learned? Biochemical Engineering Journal, 140, 57–71. https://doi.org/10.1016/j.bej.2018.09.001

Ho, P., Westerwalbesloh, C., Kaganovitch, E., Grünberger, A., Neubauer, P., Kohlheyer, D., & Lieres, E. von (2019). Reproduction of Large-Scale Bioreactor Conditions on Microfluidic Chips. Microorganisms, 7(4). https://doi.org/10.3390/microorganisms7040105

Junker, B. H. (2004). Scale-up methodologies for Escherichia coli and yeast fermentation processes. Journal of Bioscience and Bioengineering, 97(6), 347–364. https://doi.org/10.1016/S1389-1723(04)70218-2

Kaiser, M., Jug, F., Julou, T., Deshpande, S., Pfohl, T., Silander, O. K., Myers, G., & van Nimwegen, E. (2018). Monitoring single-cell gene regulation under dynamically controllable conditions with integrated microfluidics and software. Nature Communications, 9(1), 212. https://doi.org/10.1038/s41467-017-02505-0

Käß, F., Hariskos, I., Michel, A., Brandt, H.-J., Spann, R., Junne, S., Wiechert, W., Neubauer, P., & Oldiges, M. (2014). Assessment of robustness against dissolved oxygen/substrate oscillations for C. glutamicum DM1933 in two-compartment bioreactor. Bioprocess and Biosystems Engineering, 37(6), 1151–1162. https://doi.org/10.1007/s00449-013-1086-0

Käß, F., Junne, S., Neubauer, P., Wiechert, W., & Oldiges, M. (2014). Process inhomogeneity leads to rapid side product turnover in cultivation of Corynebacterium glutamicum. Microbial Cell Factories, 13, 6. https://doi.org/10.1186/1475-2859-13-6

Klask, C.-M., Kliem-Kuster, N., Molitor, B., & Angenent, L. T. (2020). Nitrate Feed Improves Growth and Ethanol Production of Clostridium ljungdahlii With CO_2_ and H_2_, but Results in Stochastic Inhibition Events. Frontiers in Microbiology, 11, 724. https://doi.org/10.3389/fmicb.2020.00724

Kovárová-Kovar, K., & Egli, T. (1998). Growth Kinetics of Suspended Microbial Cells: From Single-Substrate-Controlled Growth to Mixed-Substrate Kinetics. Microbiology and Molecular Biology Reviews, 62(3), 646–666.

Krüger, A., Wiechert, J., Gätgens, C., Polen, T., Mahr, R., & Frunzke, J. (2019). Impact of CO_2_/HCO_3_-Availability on Anaplerotic Flux in Pyruvate Dehydrogenase Complex-Deficient Corynebacterium glutamicum Strains. Journal of Bacteriology, 201(20). https://doi.org/10.1128/JB.00387-19

Langheinrich, C., & Nienow, A. W. (1999). Control of pH in large-scale, free suspension animal cell bioreactors: Alkali addition and pH excursions. Biotechnology and Bioengineering, 66(3), 171–179. https://doi.org/10.1002/(SICI)1097-0290(1999)66:3<171::AID-BIT5>3.0.CO;2-T

Lara, A. R., Galindo, E., Ramírez, O. T., & Palomares, L. A. (2006). Living With Heterogeneities in Bioreactors: Understanding the Effects of Environmental Gradients on Cells. Molecular Biotechnology, 34(3), 355–382. https://doi.org/10.1385/MB:34:3:355

Larsson, G., & Enfors, S.-O. (1988). Studies of insufficient mixing in bioreactors: Effects of limiting oxygen concentrations and short term oxygen starvation on Penicillium chrysogenum. Bioprocess Engineering, 3(3), 123–127. https://doi.org/10.1007/BF00373475

Lee, J.-Y., Na, Y.-A., Kim, E., Lee, H.-S., & Kim, P. (2016). Erratum to: The Actinobacterium Corynebacterium glutamicum, an Industrial Workhorse. Journal of Microbiology and Biotechnology, 26(7), 1341. https://doi.org/10.4014/jmb.2016.2607.1341

Lemoine, A., Limberg, M. H., Kästner, S., Oldiges, M., Neubauer, P., & Junne, S. (2016). Performance loss of Corynebacterium glutamicum cultivations under scale-down conditions using complex media. Engineering in Life Sciences, 16(7), 620–632. https://doi.org/10.1002/elsc.201500144

Lemoine, A., Maya Mart?nez-Iturralde, N., Spann, R., Neubauer, P., & Junne, S. (2015). Response of Corynebacterium glutamicum exposed to oscillating cultivation conditions in a two- and a novel three-compartment scale-down bioreactor. Biotechnology and Bioengineering, 112(6), 1220– 1231. https://doi.org/10.1002/bit.25543

Limberg, M. H., Joachim, M., Klein, B., Wiechert, W., & Oldiges, M. (2017). pH fluctuations imperil the robustness of C. glutamicum to short term oxygen limitation. Journal of Biotechnology, 259, 248– 260. https://doi.org/10.1016/j.jbiotec.2017.08.018

Limberg, M. H., Pooth, V., Wiechert, W., & Oldiges, M. (2016). Plug flow versus stirred tank reactor flow characteristics in two-compartment scale-down bioreactor: Setup-specific influence on the metabolic phenotype and bioprocess performance of Corynebacterium glutamicum. Engineering in Life Sciences, 16(7), 610–619. https://doi.org/10.1002/elsc.201500142

Lindström, S., & Andersson-Svahn, H. (2010). Overview of single-cell analyses: Microdevices and applications. Lab on a Chip, 10(24), 3363–3372. https://doi.org/10.1039/c0lc00150c

Liu, Y., Yang, X., Yin, Y., Lin, J., Chen, C., Pan, J., Si, M., & Shen, X. (2016). Mycothiol protects Corynebacterium glutamicum against acid stress via maintaining intracellular pH homeostasis, scavenging ROS, and S-mycothiolating MetE. The Journal of General and Applied Microbiology, 62(3), 144–153. https://doi.org/10.2323/jgam.2016.02.001

Lopes, M., Belo, I., & Mota, M. (2014). Over-pressurized bioreactors: Application to microbial cell cultures. Biotechnology Progress, 30(4), 767–775. https://doi.org/10.1002/btpr.1917

Mustafi, N., Grünberger, A., Mahr, R., Helfrich, S., Nöh, K., Blombach, B., Kohlheyer, D., & Frunzke, J. (2014). Application of a genetically encoded biosensor for live cell imaging of L-valine production in pyruvate dehydrogenase complex-deficient Corynebacterium glutamicum strains. PloS One, 9(1), e85731. https://doi.org/10.1371/journal.pone.0085731

Nadal-Rey, G., McClure, D. D., Kavanagh, J. M., Cornelissen, S., Fletcher, D. F., & Gernaey, K. V. (2020). Understanding gradients in industrial bioreactors. Biotechnology Advances, 107660. https://doi.org/10.1016/j.biotechadv.2020.107660

Namdev, P. K., Thompson, B. G., & Gray, M. R. (1992). Effect of feed zone in fed-batch fermentations of Saccharomyces cerevisiae. Biotechnology and Bioengineering, 40(2), 235–246. https://doi.org/10.1002/bit.260400207

Neubauer, P., & Junne, S. (2010). Scale-down simulators for metabolic analysis of large-scale bioprocesses. Current Opinion in Biotechnology, 21(1), 114–121. https://doi.org/10.1016/j.copbio.2010.02.001

Neubauer, P., & Junne, S. (2016). Scale-Up and Scale-Down Methodologies for Bioreactors. In C.-F. Mandenius (Ed.), Bioreactors (pp. 323–354). Weinheim, Germany: Wiley-VCH Verlag GmbH & Co. KGaA. https://doi.org/10.1002/9783527683369.ch11

Olughu, W., Nienow, A., Hewitt, C., & Rielly, C. (2020). Scale-down studies for the scale-up of a recombinant Corynebacterium glutamicum fed-batch fermentation: Loss of homogeneity leads to lower levels of cadaverine production. Journal of Chemical Technology and Biotechnology, 95(3), 675–685. https://doi.org/10.1002/jctb.6248

Onyeaka, H., Nienow, A. W., & Hewitt, C. J. (2003). Further studies related to the scale-up of high cell density Escherichia coli fed-batch fermentations: The additional effect of a changing microenvironment when using aqueous ammonia to control pH. Biotechnology and Bioengineering, 84(4), 474–484. https://doi.org/10.1002/bit.10805

Pohlscheidt, M., Charaniya, S., Bork, C., Jenzsch, M., Noetzel, T. L., & Luebbert, A. (2009). Bioprocess and Fermentation Monitoring. In M. C. Flickinger (Ed.), Encyclopedia of Industrial Biotechnology. Hoboken, NJ, USA: John Wiley & Sons, Inc. https://doi.org/10.1002/9780470054581.eib606.pub2

Probst, C., Grünberger, A., Braun, N., Helfrich, S., Nöh, K., Wiechert, W., & Kohlheyer, D. (2015). Rapid inoculation of single bacteria into parallel picoliter fermentation chambers. Analytical Methods, 7(1), 91–98. https://doi.org/10.1039/C4AY02257B

Reuss, M., Schmalzriedt, S., & Jenne, M. (Eds.) (1994). Structured Modelling of Bioreactors (Vol. 15). https://doi.org/10.1007/978-94-017-0641-4_29

Sahm, H., Antranikian, G., Stahmann, K.-P., & Takors, R. (Eds.) (2013). Industrielle Mikrobiologie. Berlin: Springer Spektrum.

Schindelin, J., Arganda-Carreras, I., Frise, E., Kaynig, V., Longair, M., Pietzsch, T., Preibisch, S., Rueden, C., Saalfeld, S., Schmid, B., Tinevez, J.-Y., White, D. J., Hartenstein, V., Eliceiri, K., Tomancak, P., & Cardona, A. (2012). Fiji: An open-source platform for biological-image analysis. Nature Methods, 9(7), 676–682. https://doi.org/10.1038/nmeth.2019

Spann, R., Glibstrup, J., Pellicer-Alborch, K., Junne, S., Neubauer, P., Roca, C., Kold, D., Lantz, A. E., Sin, G., Gernaey, K. V., & Krühne, U. (2019). CFD predicted pH gradients in lactic acid bacteria cultivations. Biotechnology and Bioengineering, 116(4), 769–780. https://doi.org/10.1002/bit.26868

Takors, R. (2012). Scale-up of microbial processes: Impacts, tools and open questions. Journal of Biotechnology, 160(1-2), 3–9. https://doi.org/10.1016/j.jbiotec.2011.12.010

Täuber, S., Blöbaum, L., Wendisch, V. F., & Grünberger, A. (2021). Growth Response and Recovery of Corynebacterium glutamicum Colonies on Single-Cell Level Upon Defined pH Stress Pulses. Frontiers in Microbiology, 12, 711893. https://doi.org/10.3389/fmicb.2021.711893

Täuber, S., Golze, C., Ho, P., Lieres E. von, & Grünberger, A. (2020). dMSCC: A microfluidic platform for microbial single-cell cultivation of Corynebacterium glutamicum under dynamic environmental medium conditions. Lab on a Chip. Advance online publication. https://doi.org/10.1039/D0LC00711K

Täuber, S., Lieres, E., & Grünberger, A. (2020). Dynamic Environmental Control in Microfluidic Single-Cell Cultivations: From Concepts to Applications. Small, 1906670. https://doi.org/10.1002/smll.201906670

Templer, R. H., & Ces, O. (2008). New frontiers in single-cell analysis. Journal of the Royal Society, Interface, 5 Suppl 2, S111–2. https://doi.org/10.1098/rsif.2008.0279.focus

Uhlendorf, J., Miermont, A., Delaveau, T., Charvin, G., Fages, F., Bottani, S., Batt, G., & Hersen, P. (2012). Long-term model predictive control of gene expression at the population and single-cell levels. Proceedings of the National Academy of Sciences of the United States of America, 109(35), 14271–14276. https://doi.org/10.1073/pnas.1206810109

Unthan, S., Grünberger, A., van Ooyen, J., Gätgens, J., Heinrich, J., Paczia, N., Wiechert, W., Kohlheyer, D., & Noack, S. (2014). Beyond growth rate 0.6: What drives Corynebacterium glutamicum to higher growth rates in defined medium. Biotechnology and Bioengineering, 111(2), 359–371. https://doi.org/10.1002/bit.25103

Vrábel, P., van der Lans, R. G., Luyben, K. C., Boon, L., & Nienow, A. W. (2000). Mixing in large-scale vessels stirred with multiple radial or radial and axial up-pumping impellers: modelling and measurements. Chemical Engineering Science, 55(23), 5881–5896. https://doi.org/10.1016/S0009-2509(00)00175-5

Wang, P., Robert, L., Pelletier, J., Dang, W. L., Taddei, F., Wright, A., & Jun, S. (2010). Robust growth of Escherichia coli. Current Biology, 20(12), 1099–1103. https://doi.org/10.1016/j.cub.2010.04.045.

Wehrs, M., Tanjore, D., Eng, T., Lievense, J., Pray, T. R., & Mukhopadhyay, A. (2019). Engineering Robust Production Microbes for Large-Scale Cultivation. Trends in Microbiology, 27(6), 524–537. https://doi.org/10.1016/j.tim.2019.01.006

Westerwalbesloh, C., Grünberger, A., Stute, B., Weber, S., Wiechert, W., Kohlheyer, D., & Lieres, E. von (2015). Modeling and CFD simulation of nutrient distribution in picoliter bioreactors for bacterial growth studies on single-cell level. Lab on a Chip, 15(21), 4177–4186. https://doi.org/10.1039/c5lc00646e

Wood, J. M., Bremer, E., Csonka, L. N., Kraemer, R., Poolman, B., van der Heide, T., & Smith, L. T. (2001). Osmosensing and osmoregulatory compatible solute accumulation by bacteria. Comparative Biochemistry and Physiology Part a: Molecular & Integrative Physiology, 130(3), 437– 460. https://doi.org/10.1016/S1095-6433(01)00442-1

Zimmermann, H. F., Anderlei, T., Büchs, J., & Binder, M. (2006). Oxygen limitation is a pitfall during screening for industrial strains. Applied Microbiology and Biotechnology, 72(6), 1157–1160. https://doi.org/10.1007/s00253-006-0414-6

