## Supplement information for "Microfluidic single-cell scale-down bioreactors: A proof-of-concept for the growth of *Corynebacterium glutamicum* at oscillating pH values"

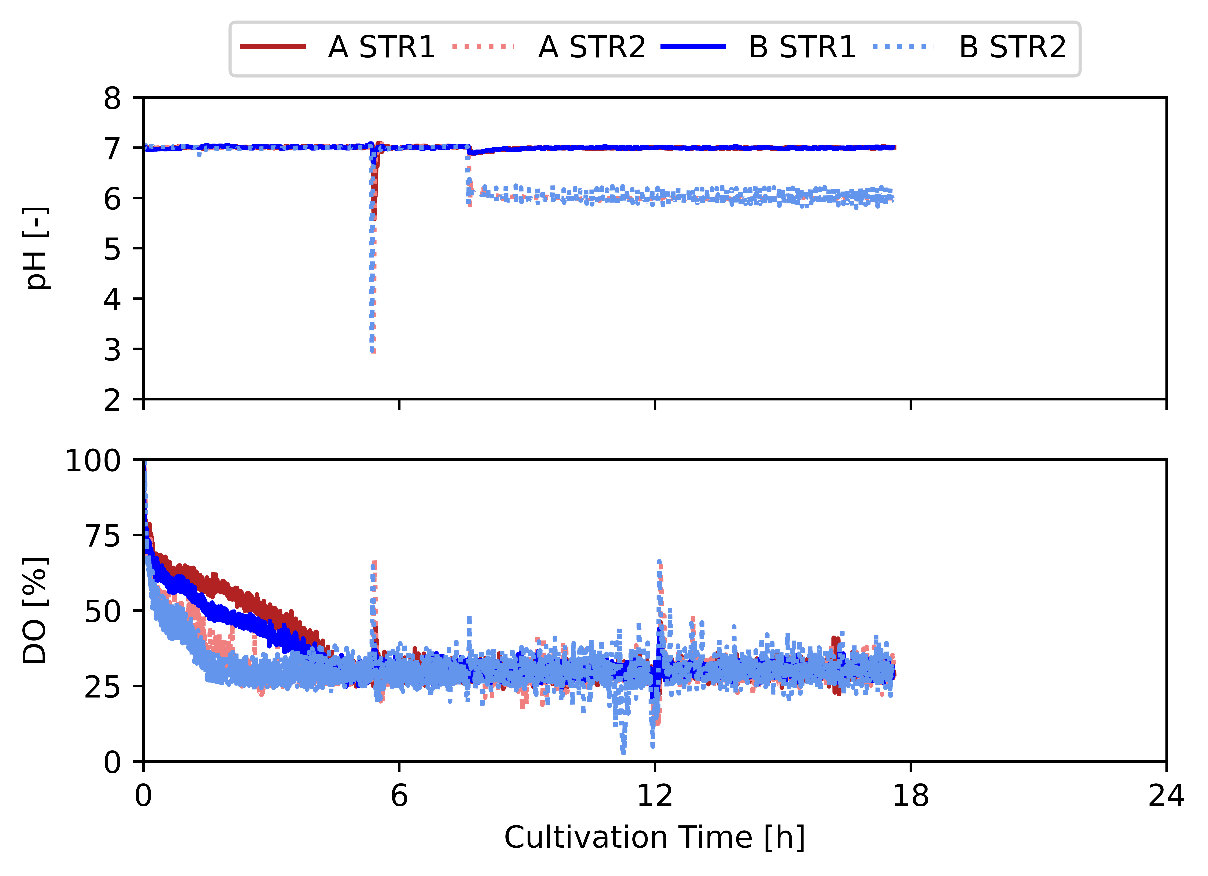

**Figure S1**: Course of the pH value and DO concentration over the process time of the two-CR cultivation of *C. glutamicum* during pH 6/7 oscillation.

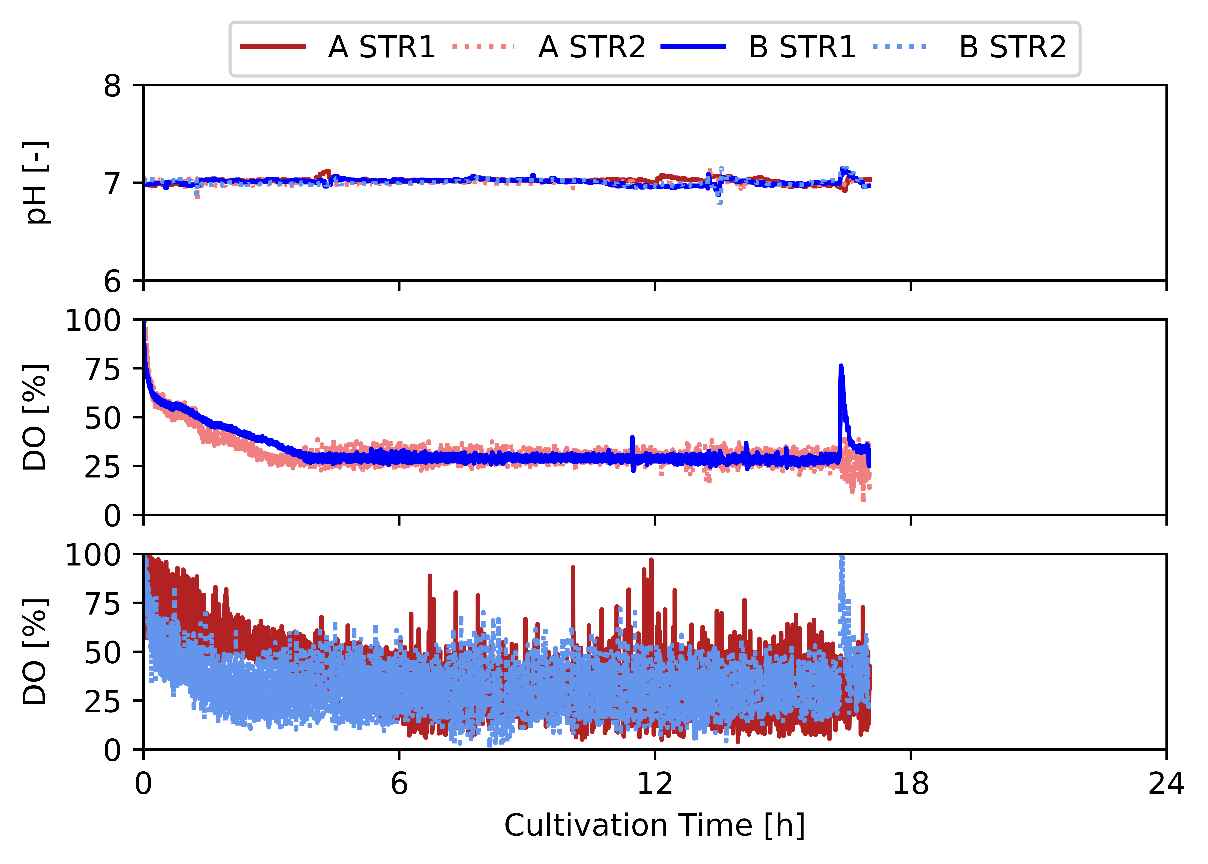

**Figure S2**: Course of the pH value and DO concentration over the process time of the two-CR cultivation of *C. glutamicum* during pH 7/7 oscillation.

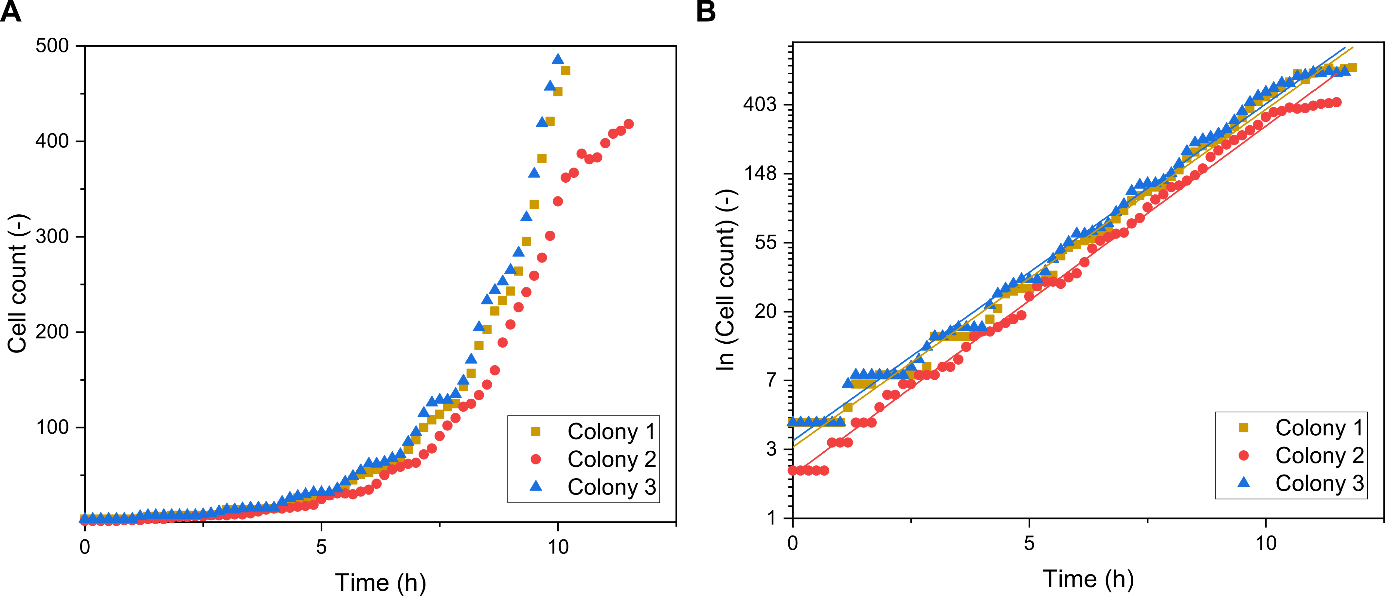

**Figure S3**: As an example of cultivation curves for pH 6 with a relative oscillation ratio *w_rel_* = 0.5 and a total interval duration of T = 20 min. A) Exponential growth. B) Semilogarithmic plot of cell count with the linear regression to calculate the growth rate. Three microcolonies are shown in red, blue and orange.

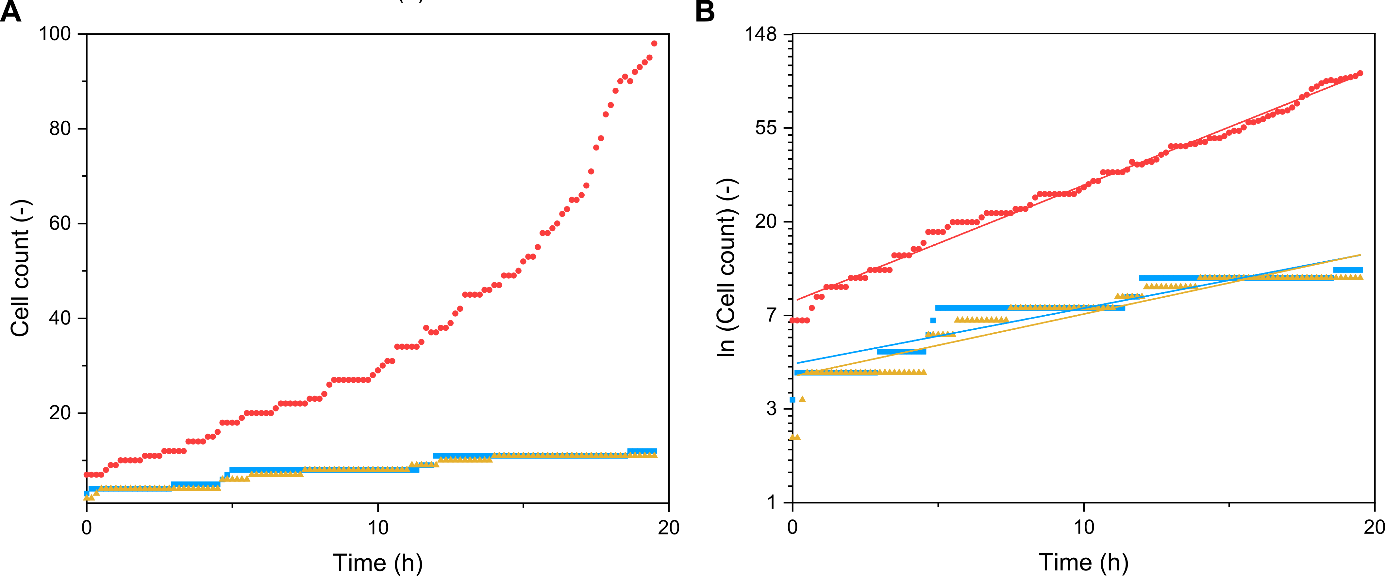

**Figure S4**: Growth curve of *C. glutamicum* during a pH 5/7 oscillation with an oscillation duration *w* = 3 min. A) Exponential growth. B) Semilogarithmic plot of cell count with the linear regression to calculate the growth rate. Three microcolonies are shown in red, blue and orange.

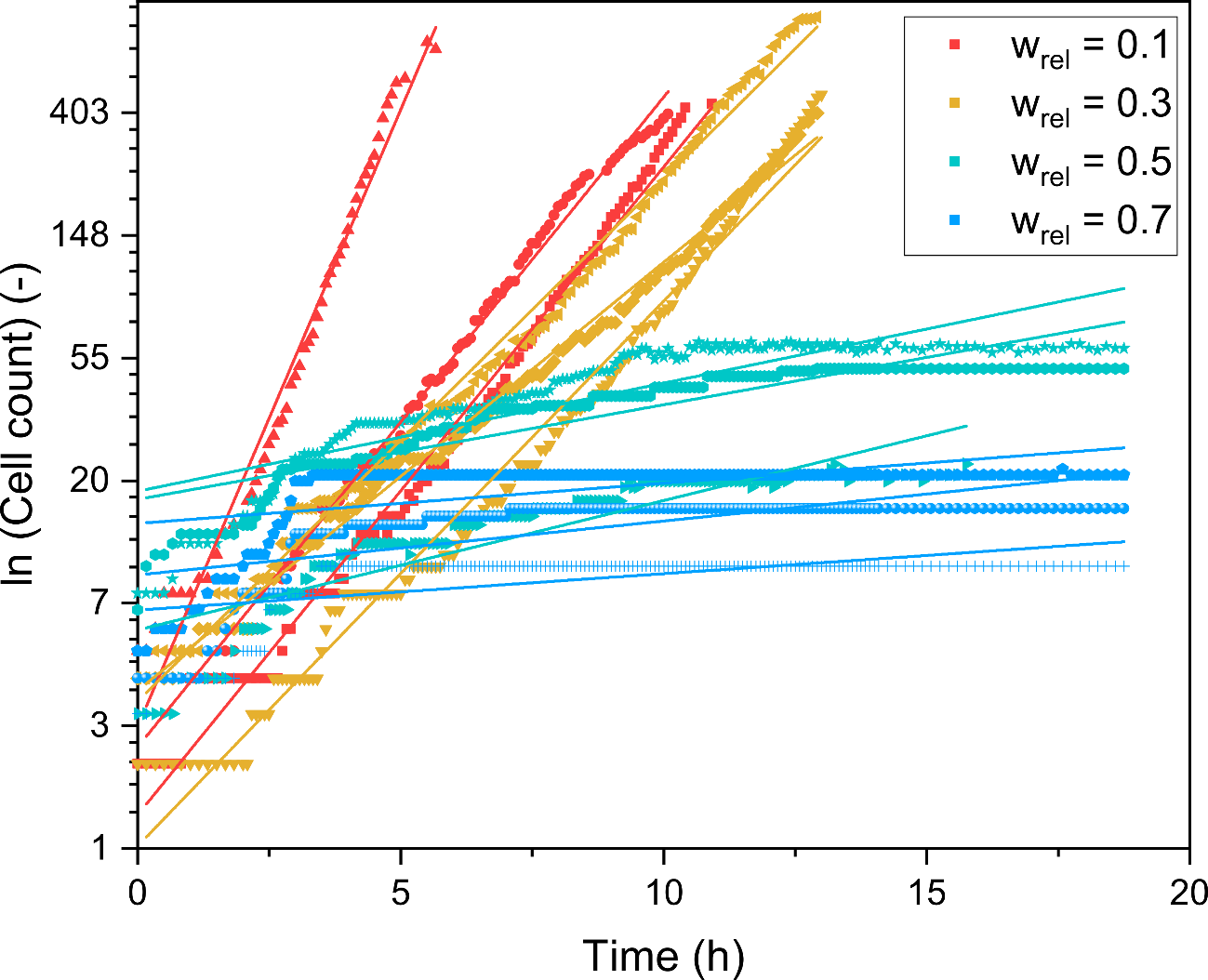

**Figure S5**: Semi-logarithmic plot of the cell count with the linear regression tocalculate growht rate of *C. glutamicum* during a pH 5/7 oscillation with a total interval duration T= 10 min. For each realtive oscillation ratio w_rel_, three microcolonies are shown in the same color.

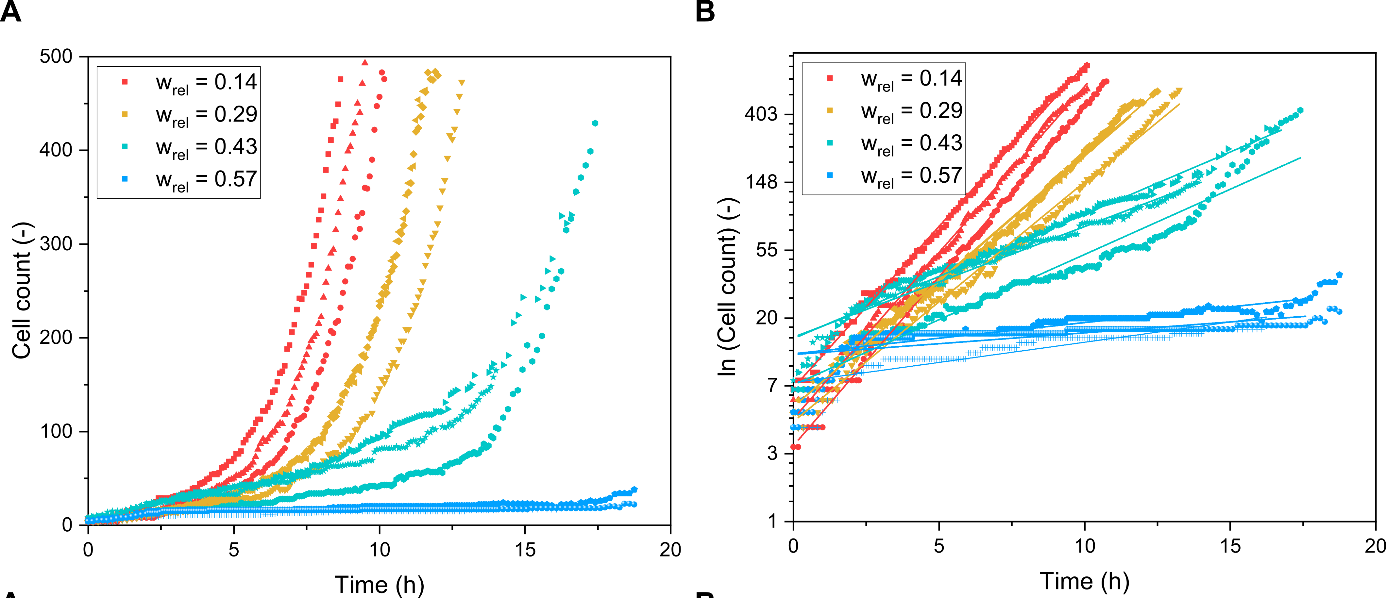

**Figure S6**: Growth curve of *C. glutamicum* during a pH 5/7 oscillation with a total interval duration T= 35 min. A) Exponential growth. B) Semilogarithmic plot of cell count with the linear regression to calculate the growth rate. For each realtive oscillation ratio w_rel_, three microcolonies are shown in the same color.

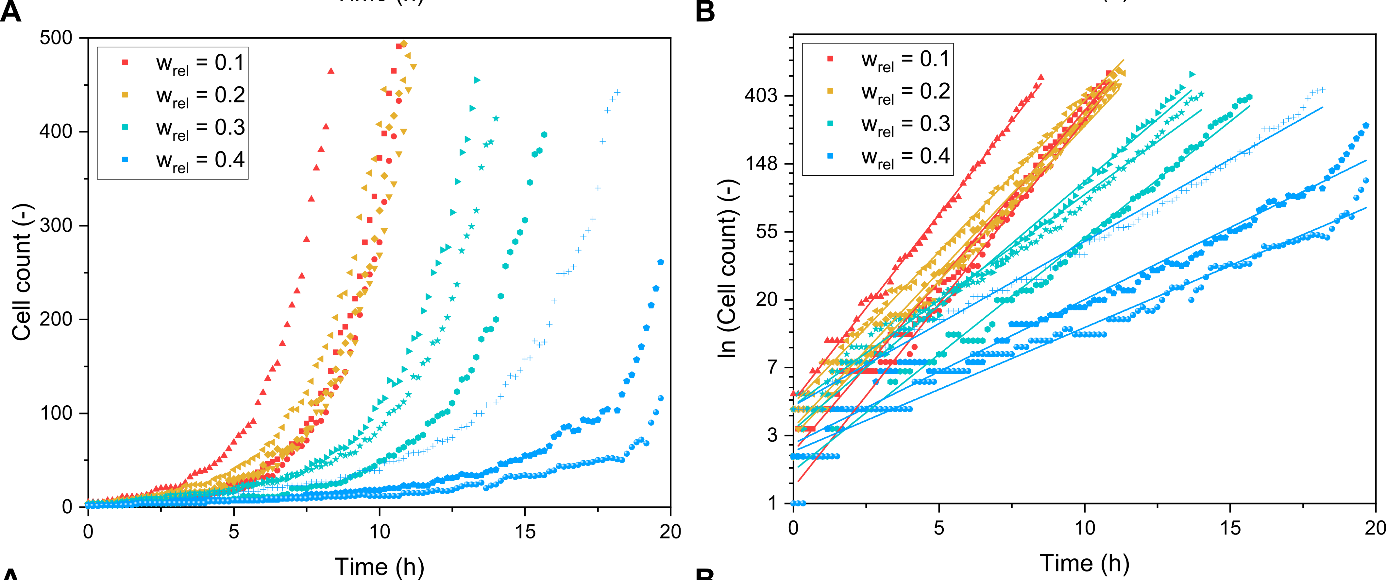

**Figure S7**: Growth curve of *C. glutamicum* during a pH 5/7 oscillation with a total interval duration T= 50 min. A) Exponential growth. B) Semilogarithmic plot of cell count with the linear regression to calculate the growth rate. For each realtive oscillation ratio w_rel_, three microcolonies are shown in the same color.

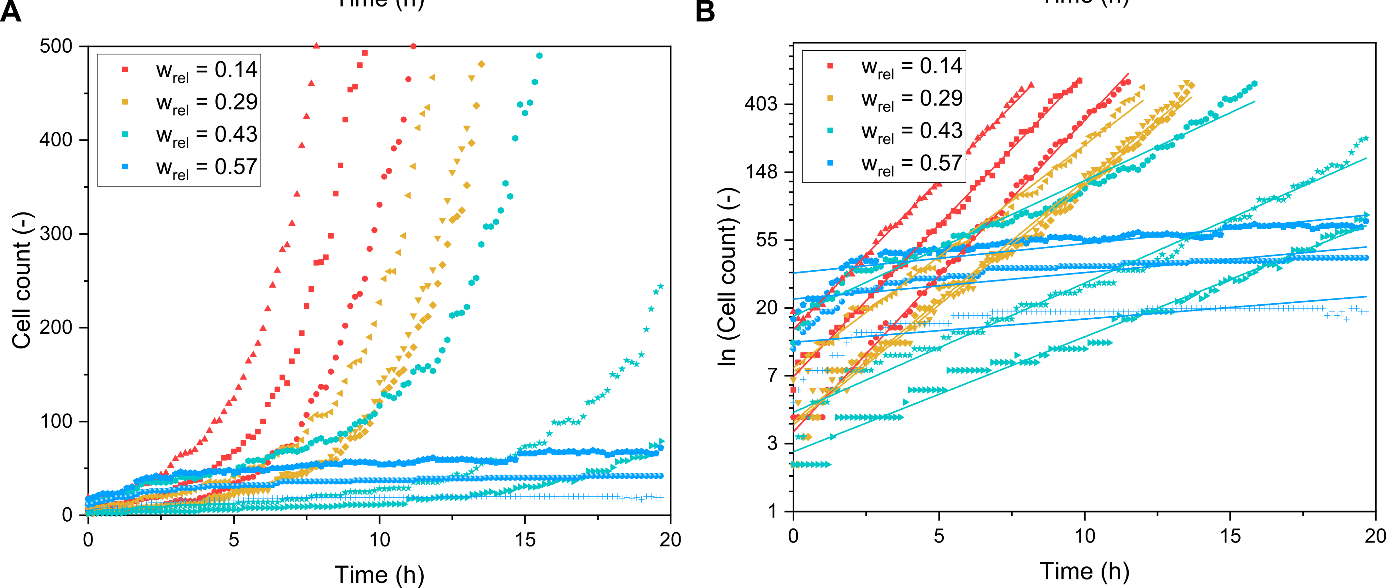

**Figure S8**: Growth curve of *C. glutamicum* during a pH 5/7 oscillation with a total interval duration T= 70 min. A) Exponential growth. B) Semilogarithmic plot of cell count with the linear regression to calculate the growth rate. For each realtive oscillation ratio w_rel_, three microcolonies are shown in the same color.

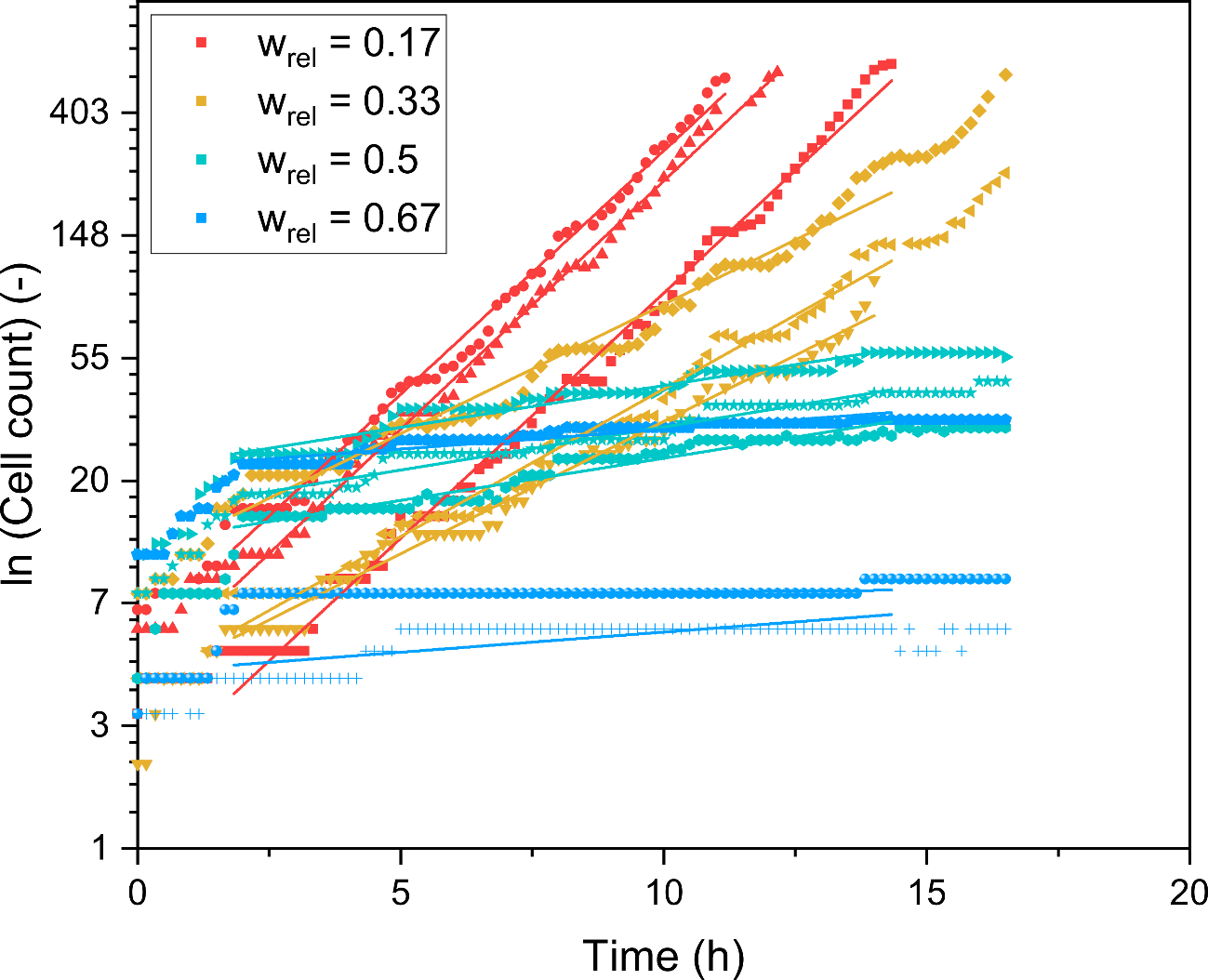

**Figure S9**: Semi-logarithmic plot of the cell count with the linear regression tocalculate growht rate of *C. glutamicum* during a pH 5/7 oscillation with a total interval duration T= 180 min. For each realtive oscillation ratio w_rel_, three microcolonies are shown in the same color.

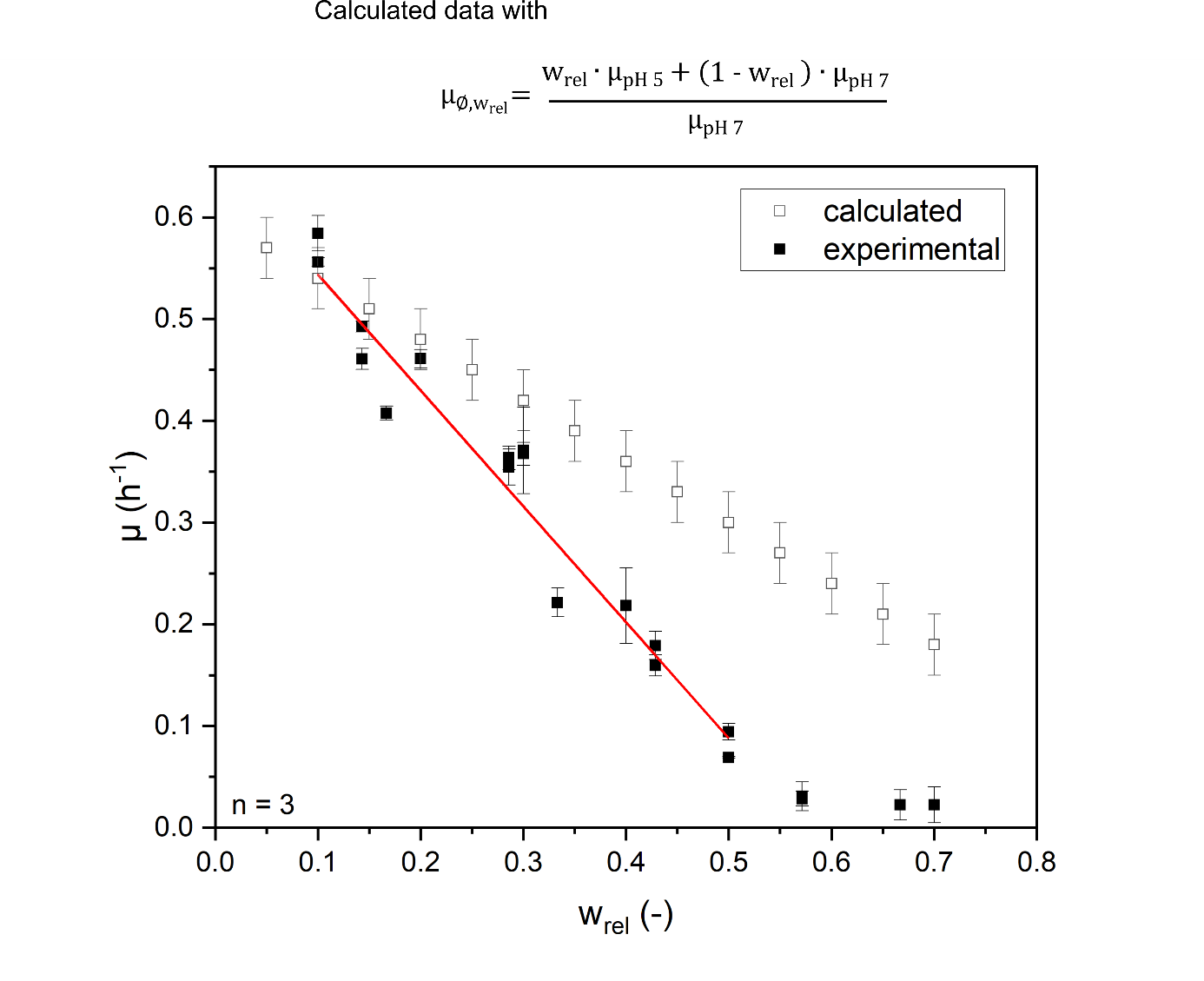

**Figure S10:** Comparison of the complied experimental data of Figure 4B and the assumed data set, if µ_pH 5_ = 0 h^-1^ and µ_pH 7_ = 0.58 ± 0.03 h^-1^.

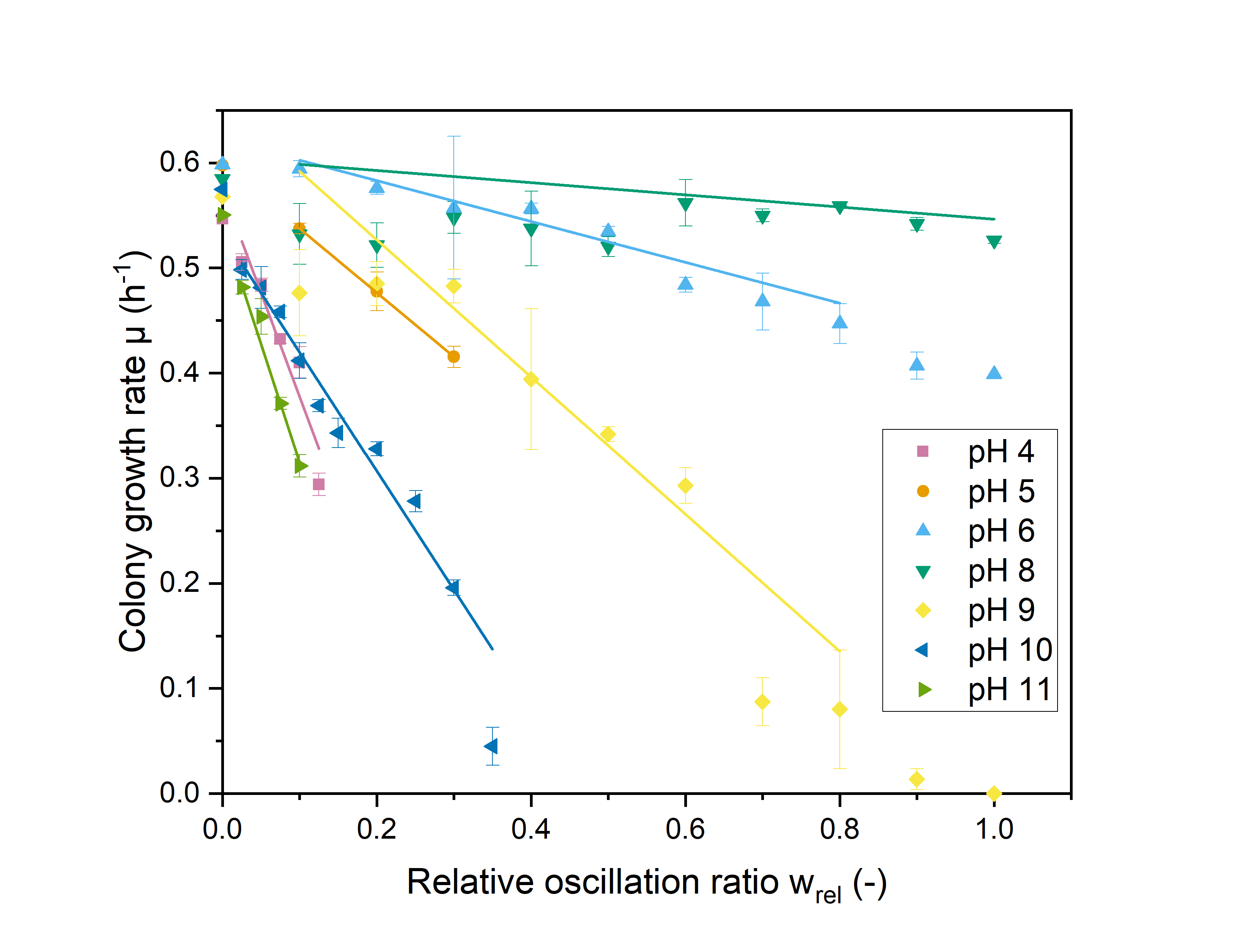

**Figure S11**: Growth rate and linear regression depended on the different relative oscillation ratios *w_rel_*. Variation of relative oscillation ratio *w_rel_* with a total interval duration *T* of 20 min for pH stress amplitudes *A* 4 - 11.
